# Targeting BTN2A1 enhances Vγ9Vδ2 T cell effector functions and triggers tumor cells pyroptosis

**DOI:** 10.1101/2023.06.15.545049

**Authors:** Anne Charlotte le Floch, Caroline Imbert, Nicolas Boucherit, Laurent Gorvel, Stéphane Fattori, Florence Orlanducci, Aude Le Roy, Lorenzo Archetti, Lydie Crescence, Laurence Panicot-Dubois, Christophe Dubois, Norbert Vey, Antoine Briantais, Amandine Anastasio, Carla E. Cano, Geoffrey Guittard, Mathieu Frechin, Daniel Olive

## Abstract

Vγ9Vδ2 T cells are potent but elusive cytotoxic effectors. Means to stimulate their function could lead to powerful new cancer immunotherapies. BTN2A1, a surface protein has recently been shown to bind the Vγ9 chain of the γδ TCR but its precise role in modulating Vγ9Vδ2 T cells functions remains unknown.

Here we show that 107G3B5, a monoclonal anti-BTN2A1 agonist antibody, significantly enhances Vγ9Vδ2 T cell functions against hematological or solid cell lines and against primary cells from adult acute lymphoblastic leukemia patients. New computer vision strategies applied to holotomographic microscopy videos show that 107G3B5 enhances the interaction between Vγ9Vδ2 T cells and target cells in a quantitative and qualitative manner. In addition, we provide evidence that Vγ9Vδ2 T cells activated by 107G3B5 induce caspase 3/7 activation in tumor cells, thereby triggering their death by pyroptosis.

We thus demonstrate that targeting BTN2A1 with 107G3B5 enhances the Vγ9Vδ2 T cell antitumor response by triggering the pyroptosis-induced immunogenic cell death.

## Introduction

In contrast to αβ T cells, γδ T cell activation is MHC-unrestricted and relies on the detection of various host cell-derived molecules^1^ or exogenous pathogens by both TCR and non-TCR receptors. γδ T cells represent the earliest source of IFNγ secretion in the tumor microenvironment^2^ and recent transcriptome analyses of human tumors reveal that high γδ T infiltration has the best prognostic value in comparison to other immune subsets^3, 4^. Vγ9Vδ2 T cells are the major subtype of blood γδ T cells and are activated by non-peptidic phosphorylated metabolites, called phosphoantigens (pAgs). These pAgs are produced by transformed or infected cells^5, 6^ via the human mevalonate pathway during the biosynthesis of isoprenoids^7^.

A better understanding of the elusive mechanisms of action of Vγ9Vδ2 T cells will lead to new potential immunotherapies. In this context, the interaction between BTN3A and BTN2A1 at the plasma membrane, which enables the recognition of phosphoantigens (pAgs) by Vγ9Vδ2 T cells through an “inside-outside signaling” ^8–10^ is of particular interest. BTN3A1 cooperates with BTN2A1 for cell surface export^27^ and binds to phosphoantigens (pAgs) through its intracellular B30.2 domain which also stabilizes its interaction with BTN2A1^11, 12^ with pAgs are acting as a “glue”. Most importantly, BTN2A1 is the direct ligand for the Vγ9 chain of the γδ TCR^13, 14^, representing a fundamental new paradigm in Vγ9Vδ2 T cell biology. While it is clear that BTN2A1 is mandatory for Vγ9Vδ2 T cell activation in tumoral context^10, 13, 14^, its precise role in modulating the functions of Vγ9Vδ2 T cells needs to be carefully investigated.

Cytotoxic functions of Vγ9Vδ2 T cells have been demonstrated in preclinical models of solid cancers^15–19^ and myeloid or lymphoid hematologic malignancies^20–23^. Therefore, potential immunotherapies based on Vγ9Vδ2 T cells have been rapidly evaluated in clinical trials, including in cancer patients with advanced solid or hematological cancers. Most studies used pharmacological inducers of pAgs, such as aminobisphosphonates^24^. While these studies tended to show objective responses, they were associated with limited efficacy results^25^.

Low tumor immunogenicity and a cold tumor microenvironment are two of the main features that occur in different cancer types and are responsible for resistance to immunotherapy. One possibility is to transform ‘cold’ tumors into ‘hot’ tumors in particular by inducing immunogenic tumor cell death ^26^. Natural killer cells and cytotoxic T cells were thought to induce tumor cell death mainly by non-inflammatory apoptosis, however, several studies have recently revealed that these cells can suppress tumors by inducing immunogenic cell death including pyroptosis ^27, 28^. GSDME (Gasdermin E) is activated by inflammatory caspases 3/7 and can form oligomers that insert into membranes to form pores, thereby mediating pyroptotic cell death. The Induction of non-apoptotic cell death has attracted much attention in the field of antitumor therapy, but until now the ability of Vγ9Vδ2 T cells to induce pyroptosis or other immunogenic cell death remained unknown.

Here, we show that a unique agonist mAb 107G3B5 targeting BTN2A1 enhances the cytotoxic activity of Vγ9Vδ2 T cells. We used several *in vitro* tumor models, including ALL in both allogeneic and autologous settings, as well as solid tumor models. Newly developed computer vision powered by artificial intelligence was used to analyze videos recorded with label-free holotomographic microscopy this to demonstrate that 107G3B5 increased quantity and quality of Vγ9Vδ2 T cell interactions with target cells. These cellular events were associated with a clustering of BTN2A1 and its colocalization with BTN3A1, as demonstrated by confocal microscopy and FRET experiments. Finally, we demonstrated that activation of Vγ9Vδ2 T cells using 107G3B7 induced pyroptosis, an immunogenic cell death of cancer cells.

## Results

### BTN2A1 targeting with 107G3B5 markedly strengthens Vγ9Vδ2 T cell killing and cytokine production against many cancer cell lines

First the expression of BTN2A was investigated at the transcriptomic level in normal and tumor tissues (Supplementary Fig. 1a and b). BTN2A1 is ubiquitously expressed with a great heterogeneity across tissues or tumor subtypes. BTN2A1 also shows a distinct expression pattern compared to BTN2A2 and BTN3A1. Using publicly available scNA-Seq datasets of tissues of normal or tumor origin, we observed that BTN2A1 is ubiquitously expressed in immune and non-immune cells (Supplementary Fig. 1c). We next assessed BTN2A1 levels at the plasma membrane of tumor cells, using an anti-BTN2A 7.48 antagonist mAb and 107G3B7 mAb (Supplementary Fig.2).

Given the recently described key role of BTN2A1 in the activation of Vγ9Vδ2 T cells ^10, 13^, we hypothesized that the antibody-mediated agonistic activity of anti-BTN2A1 may enhance the cytotoxic functions of Vγ9Vδ2 T cells. To test this hypothesis, we generated monoclonal antibodies against the ectodomain of BTN2A1 by mouse immunization. Hybridoma culture supernatants displaying the highest affinity for BTN2A1 underwent two rounds of selection for their ability to enhance Vγ9Vδ2 T cell degranulation against Daudi cells (Fig. 1a). Out of 71 clones, only four clones increased the proportion of CD107ab+ Vγ9Vδ2 T cells compared to control isotype. However, sequencing of the VH/VL fragments revealed only one common unique sequence among these four clones (data not shown). The anti-BTN2A1 encoded by this sequence was named 107G3B5. We then sought to further characterize the agonistic activity of 107G3B5 on Vγ9Vδ2 T cell cytotoxicity.

**Fig. 1.**
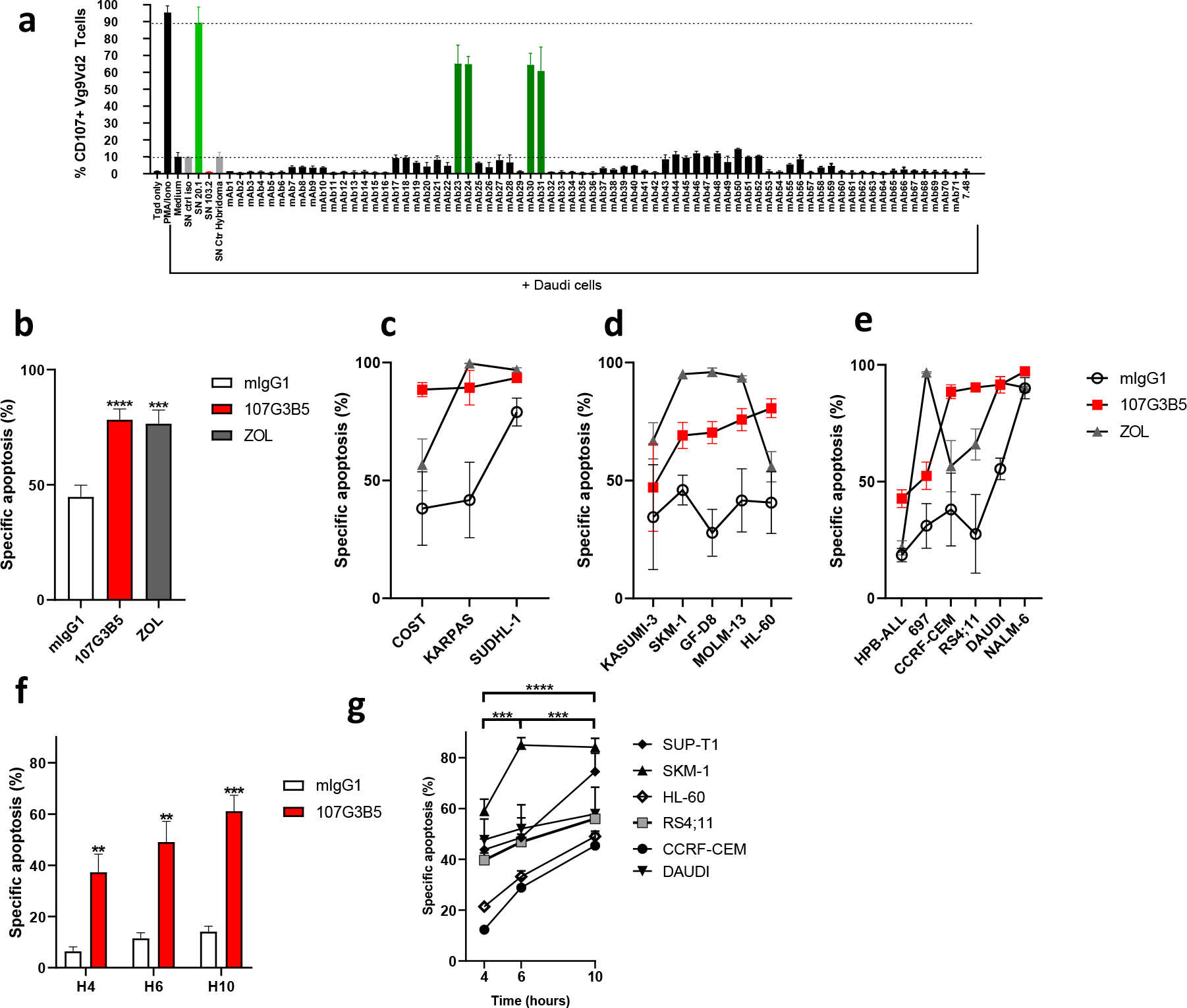
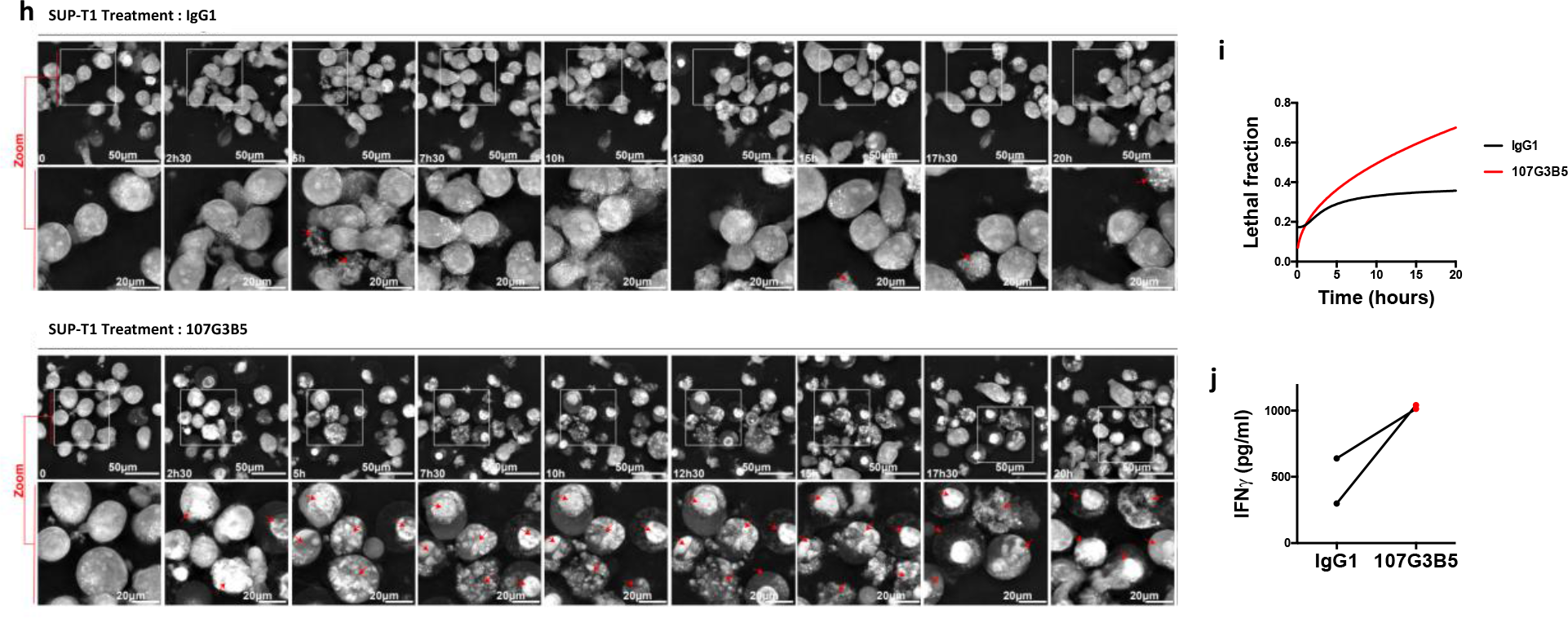
Anti-BTN2A1 107G3B5 enhances Vγ9Vδ2 T cell-mediated killing of a broad range of hematological cancer cell lines. (**a**) Vg9Vd2 T cells expanded from healthy donor PBMCs (n=3) were co-cultured with Daudi cells (E: T ratio 1:1) in the presence of 50 µL of the indicated anti-BTN2A hybridoma supernatants and %CD107ab+ cells were assessed after 4h by flow cytometry. (**b**) Apoptosis of 15 hematological cancer cell lines, including 3 ALCL cell lines (**c**), 7 ALL and NHL cell lines (**d**), and 5 AML cell lines (**e**) was evaluated by flow cytometry analysis of caspase 3/7 cleavage after 4-hour coculture with Vγ9Vδ2 T cells from 3 HV (E:T ratio 5:1) in the presence of an isotype control or anti-BTN2A 107G3B5 mAb (10µg/mL). Target cells preincubated over-night (O/N) with zoledronate (45 µM) were used as positive control. (**f,g**) Apoptosis of 6 hematological cancer cell lines was monitored by flow cytometry analysis of caspase 3/7 cleavage after 4, 6 and 10-hours of coculture with Vγ9Vδ2 T cells from 3 HV (E:T ratio 2:1) in the presence of an isotype control or anti-BTN2A 107G3B5 mAb (10µg/mL). (**h**) Sequence of snapshots depicting γδ T cells (E) coculture with SUPT1 (T; ratio T/E = 1/2) treated with 10ug/ml of mIgG1 control isotype (upper panel) or anti-BTN2A1 107G3B5 (down panel) within 20hours. The red arrows indicate apoptotic cells. (**i**) Quantification of lethal fraction. (**j**) IFNγ concentration in the culture supernatants at the end of the acquisition. Means ± s.e.m. pooled from one experiment (**b,c,d,e**) or two experiments (**f,g**) are depicted. Statistical significance was established using one-way Anova with Tukey’s post-test (**b**), Friedman test with Dunn’s post-test (**c,d,e**) or two-way Anova with Tukey’s post-test (**f,g**),. *p< 0.05; **p < 0.01; ***p < 0.001; ****p < 0.0001

As previously described^10^, the relative abundance of BTN2A1 on cell lines revealed a wide distribution in both hematological cell lines (Supplementary Fig. 2a,b) and solid cancer cell lines (Supplementary Fig. 2c). We showed that BTN2A1 was also found at the membrane of PB blasts from 28 ALL patients at diagnosis (Supplementary Fig. 2c). Collectively, these data demonstrate that BTN2A1 is ubiquitously expressed at the transcript and protein level, and that BTN2A1 is found on the cell surface of different tumor origins, including primary tumor cells.

Apoptosis assays of hematological cancer cell lines were performed after 4-hours of coculture with Vγ9Vδ2 T cells expanded from 3 HV. This revealed that 107G3B5 and zoledronate (ZOL) pretreatment significantly increased Vγ9Vδ2 T cell-mediated apoptosis to a similar extent (Fig. 1b). Different susceptibility profiles were found in anaplastic large cell lymphoma cell lines (Fig. 1c), ALL, non-Hodgkin lymphoma cell lines (Fig. 1d) and acute myeloid cell lines (Fig. 1e). An increased activity of 107G3B5 was observed over time, indeed the killing of 6 hematological cancer cell lines was higher after 6 and 10 hrs of co-culture (Fig. 1f and 1g). In addition, 107G3B5 improved both degranulation and production of TNFα and IFNγ by Vγ9Vδ2 T cells after 4 hours of co-culture with hematological cell lines (Supplementary Fig.3a and b).

To better describe the effects of 107G3B5 over time, holotomographic microscopy was used. This measures the spatial distribution of the refractive index (RI) of the observed biological object, here Vγ9Vδ2 T cells, which are small and dense in dry mass. As RI and dry mass content are linearly related, Vγ9Vδ2 T cells were the brightest on the RI image. Cancer cells were larger, had a lower dry mass density and therefore appeared darker on the RI image. Using Artificial Intelligence (AI) for signal processing with advanced thresholding methods, the RI signal alone was used for cell segmentation (Supplementary Fig. 4)^29^. This method has the significant advantage to be label-free and is a valid approach to further characterize well-known apoptotic features such as: cell budding, chromatin condensation, micronuclei, and apoptotic bodies^30^. We used SUP-T1 cells, a T-ALL model, which is representative of the average response to 107G3B5 treatment regarding Vγ9Vδ2 T cells activation and cytotoxicity (Supplementary Fig.3a and b and Fig.1g). Indeed, SUP-T1 cells co-cultured with Vγ9Vδ2 T cells showed distinguishable Vγ9Vδ2 T cells activation leading to SUP-T1 killing upon 107G3B5 treatment. In parallel almost no apoptotic events were observed in the control condition (Fig. 1h, Supplementary Movie 1 and 2). We also confirmed our results by demonstrating an increase of the lethal fraction over time (Fig. 1i). This was associated with an increased production of IFNγ (Fig. 1j). Thus, 107G3B5 improved the killing of a wide range of hematological cell lines using a holotomographic microscopy setting. This broad effect is probably due to the ubiquitous expression of BTN2A1 on various type of tumor cells (Supplementary Fig.2)^10^, which allows Vγ9Vδ2 T enhanced degranulation and Th1 cytokine production upon 107G3B5 treatment.

### 107G3B5 improves allogeneic and autologous Vγ9Vδ2 T cell functions against primary lymphoblastic blasts

The anti-tumor functions of Vγ9Vδ2 T cells have been extensively studied in a variety range of hematological malignancies ^22, 23, 31–33^ but have been less explored in ALL^20, 34–36^. Given the promising results obtained with 107G3B5 in ALL cell lines, we next sought to investigate whether 107G3B5 may enhance the functions of allogeneic Vγ9Vδ2 T cells against primary ALL blasts. Compared to the control condition, 107G3B5 increased Vγ9Vδ2 T cell-mediated apoptosis of 17 primary ALL blasts (Fig. 2a). As previously found in hematological cell lines, 107G3B5 enhanced the degranulation and Th1 cytokine production capacity of Vγ9Vδ2 T cells in co-culture with primary ALL blasts (Fig. 2b). This agonistic effect is increasing over time (Fig. 2c, and d). Additionally, 107G3B5 treatment induced sustained IFNγ secretion by allogeneic Vγ9Vδ2 T cells co-cultured with 3 primary lymphoblasts (Fig. 2e).

**Fig. 2.**
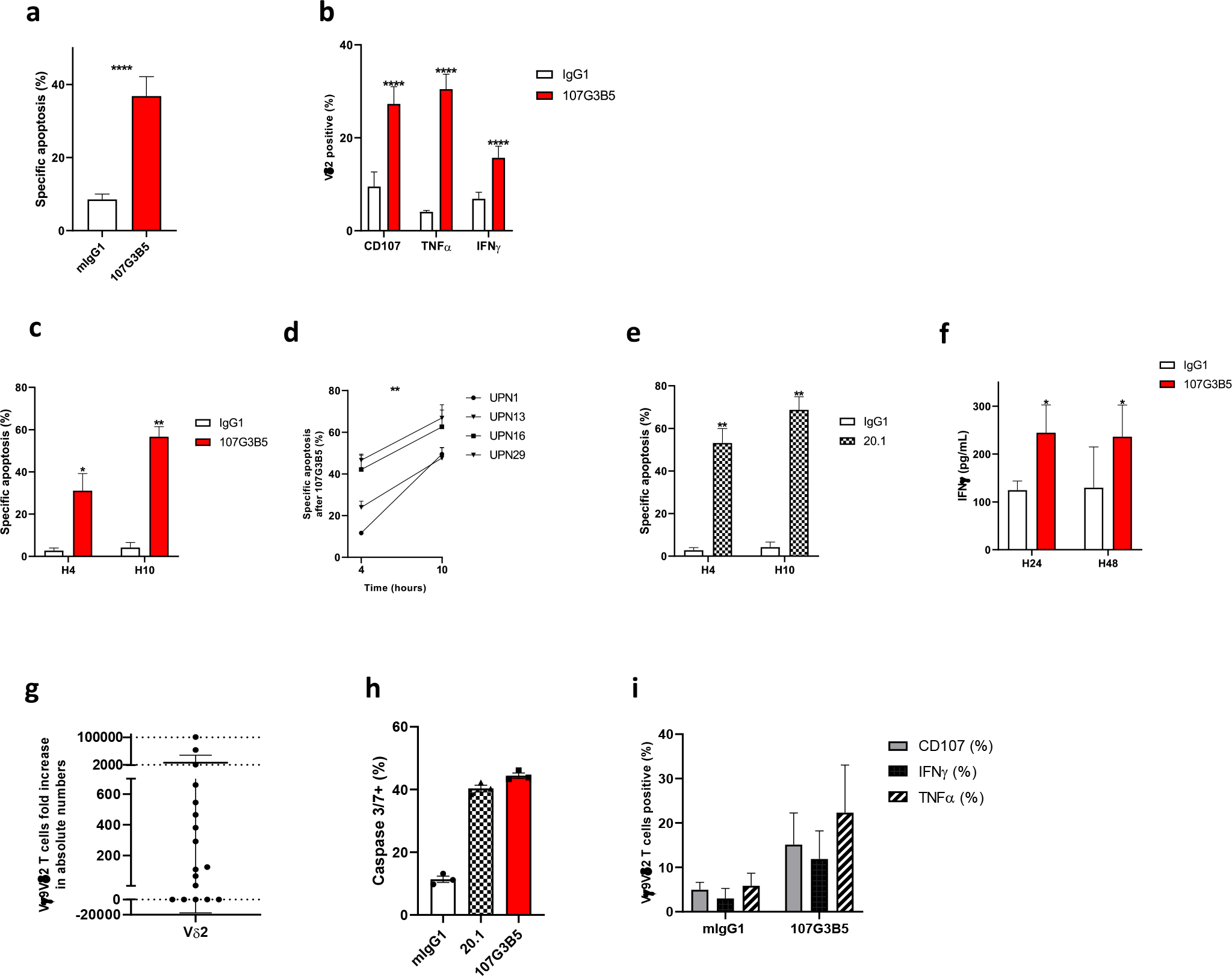
107G3B5 improves allogeneic and autologous effector response of Vγ9Vδ2 T cells against primary lymphoblastic blasts. **a** Apoptosis of 17 primary ALL blasts was assessed by analysis of caspase 3/7 cleavage after 4-hour coculture with Vγ9Vδ2 T cells from 3 HV (E:T ratio 15:1) with either isotype control or 107G3B5 (10µg/mL). (**b)** Effector functions of Vγ9Vδ2 T cells (HV, n=2 or 3) were assessed after a 4 hours co-culture with 17 primary ALL blasts (E:T ratio 1:1) in the presence of 107G3B5 or its isotype control (1µg/mL). Pooled from 6 independent experiments (**a,b,c**). (**c**-**e**) Apoptosis of 4 primary ALL blasts was monitored by analysis of caspase 3/7 cleavage after 4 and 10-hours coculture with Vγ9Vδ2 T cells from 3 HV (E:T ratio 10:1) in the presence of an isotype control, 107G3B5 or anti-BTN3A 20.1 agonist mAb (10µg/mL). (**f**) IFNγ concentration in supernatants of 24 and 48-hour coculture of Vɣ9Vδ2 T cells from 6 HV with 3 primary ALL blasts either with isotype control or 107G3B5 (10µg/mL). (**g**) PBMCs from ALL patients at diagnosis (n = 17) were cultured with ZOL for 14 days. (**h**) Apoptosis of primary ALL blasts (n=1) was evaluated by analysis of caspase 3/7 cleavage after a 4-hour coculture with autologous Vγ9Vδ2 T cells (E:T ratio 10:1), in the presence of isotype control, 107G3B5 or anti-BTN3A 20.1 agonist mAb (10µg/mL). The experiment was performed in triplicate. **i** Effector functions of Vγ9Vδ2 T cells (n=4) were assessed after a 4 hours co-culture with autologous primary ALL blasts (E:T ratio 1:1) in the presence of 107G3B5 (10µg/mL) or its isotype control. The statistical significance was established using a paired T-test (**a**), Wilcoxon test (**b**), two-way anova test (**c,d,e,f**) \**p*< 0.05; \*\**p* < 0.01; \*\*\*\**p* < 0.0001

The antileukemic response of autologous Vγ9Vδ2 T cells has been poorly studied in ALL^35^ and to our knowledge, their expansion capacities after stimulation with pAg were unknown. Using ZOL-stimulation combined with IL-2 plus IL-15 for 14 days, we expanded Vγ9Vδ2 T cells from PBMCs of 17 ALL patients. After 14 days of culture, Vγ9Vδ2 T cells from ALL patients were able to expand, with a median fold-increase of 124.8 [0.0-101 211] (Fig. 2g). In one patient, Vγ9Vδ2 T cell expansion allowed further testing of their cytotoxic capacity, which was improved after treatment of the expanded cells with 107G3B5 (Fig. 2h). As in allogeneic assays, 107G3B5 increased degranulation and cytokine secretion of Vγ9Vδ2 T cells expanded from 4 ALL patients (Fig. 2i).

Taken together, these findings consolidate the previous results obtained on cell lines, we show that Vγ9Vδ2 T cells from ALL patients proliferate in response to 107G3B5 treatment and also enhance both allogeneic and autologous Vγ9Vδ2 T cell functions against primary ALL cells.

### 107G3B5 sensitizes Vɣ9Vδ2 T cell cytotoxicity in 3D tumor spheroid models

Next, we evaluated the activity of 107G3B5 in a solid tumor setting. Using cell lines of different tumor origins, we found that 107G3B5 significantly increased Vγ9Vδ2 T cell-mediated killing compared to the control condition (Fig. 3a). To assess 107G3B5 activity in a relevant solid tumor model, we further investigated the effect of 107G3B5 on Vγ9Vδ2 T cell effector functions using 3D spheroid assays and demonstrated that 107G3B5 increased significantly Vγ9Vδ2 T cell-mediated apoptosis of Ishikawa, HT-29, CaSki and PC-3 spheroids (Fig. 3b and c).

**Fig. 3.**
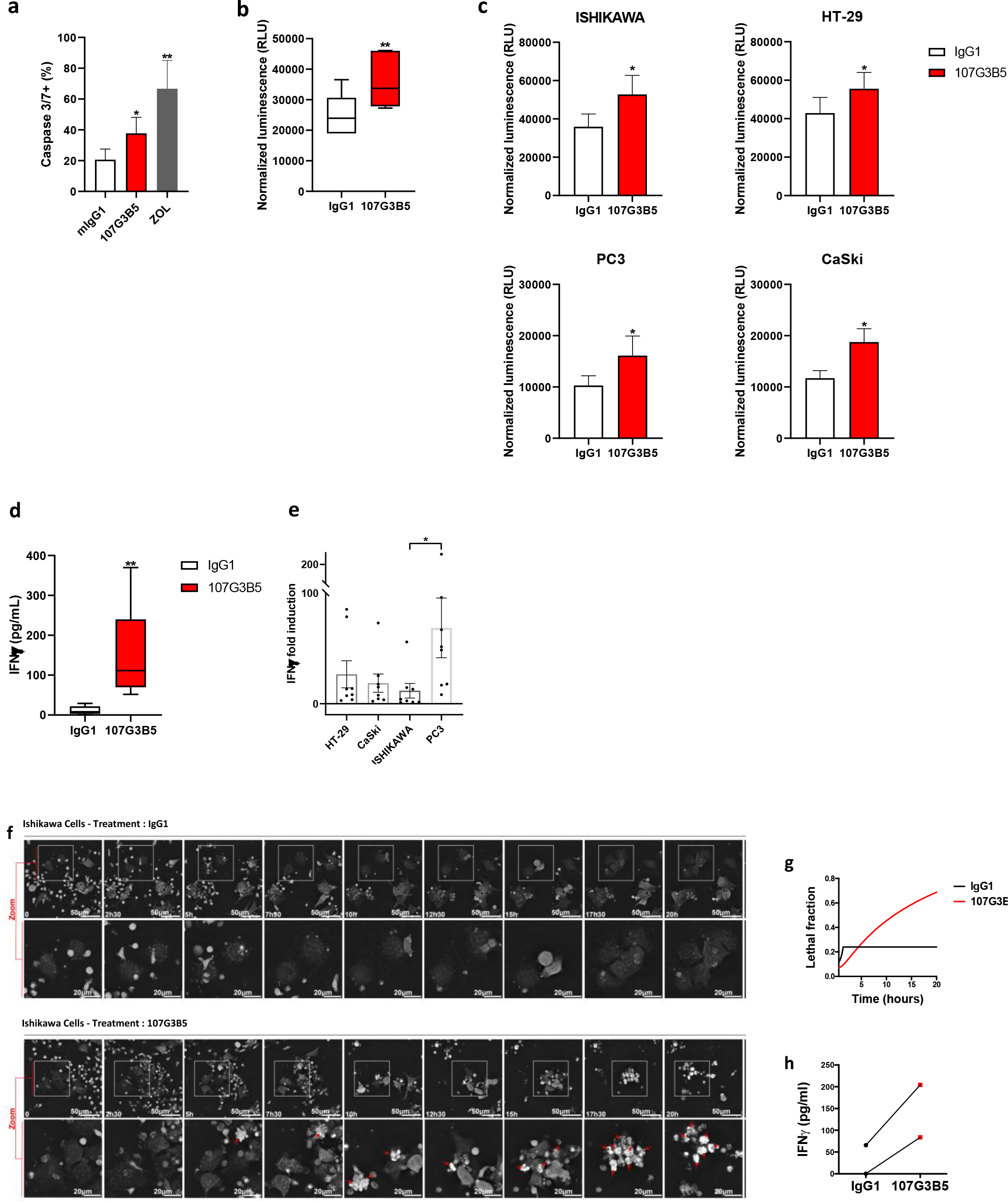
Anti BTN2A1 107G3B5 increases interactions between target cells and Vγ9Vδ2 T cells. (**a**) Apoptosis of 2D solid cancer cell lines (MCF-7, HT-29, DoTc2, PC-3, CaSki, PANC-1) was assessedby flow cytometry analysis of caspase 3/7 cleavage after 10-hour coculture with Vγ9Vδ2 T cells from 5 or 6 HV (E:T ratio 5:1) in presence of an isotype control, 107G3B5 mAb (10µg/mL). (**b, c**) Caspase 3 and 7 activity of tumor spheroids (Ishikawa, HT-29, PC-3 and CaSki) after 24-hrs coculture with Vγ9Vδ2 T cells from 6 HV (E:T ratio 5:1) in the presence of an isotype control, 107G3B5 mAb (10µg/mL). Luminescence is normalized by subtracting spontaneous caspase 3/7 activity for each spheroid. Box-plots represent mean luminescence of the four tumor spheroids (**c**). (**d,e**) IFNγ concentration in supernatants of 24-hour coculture of Vɣ9Vδ2 T cells (HV, n=8) with 3D spheroids from Ishikawa, HT-29, PC-3 and CaSki cell lines (E:T ratio 15:1) with either isotype control or 107G3B5 mAb (10µg/mL). Box-plots represent the IFNγ concentration assayed in the four tumor spheroids relative to control condition **(e)**. (**f**) Sequence of snapshots depicting γδ T cells cocultured with Ishikawa (E:T ratio 1:3) treated with 10µg/mL of mIgG1 control isotype (toppanel) or anti-BTN2A1 107G3B5 (bottompanel) during 20hours. The red arrows indicate apoptotic cells. (**g**) Quantification of the lethal fraction in HTM videos **h** IFNγ concentration in the culture supernatants at the end of the acquisition. Experiments were performed in biological (**b,c**) or technical duplicate (**d,e**).The statistical significance was established using one-way Anova with Tukey’s post-test (**a**), using Wilcoxon test (**b,c,d**) or Kruskal-Wallis test with Dunn’s post-

Live-cell fluorescence images of PC-3 spheroids co-cultured with Vγ9Vδ2 T cells from 6 HV showed a more pronounced decrease in calcein area over time after treatment with 107G3B5 compared to the isotype control condition (Supplementary Fig. 5a, b and c). After 24 hours, calcein area was significantly reduced in PC-3 spheroids after 107G3B5 treatment compared to the control condition (Supplementary Fig. 5b).

Analysis of the 24-hour coculture supernatant revealed a significant increase in IFNγ secretion after 107G3B5 treatment compared to the control condition (Fig. 3d). Consistent with previous findings, the PC-3 cell line appeared to be highly susceptible to Vγ9Vδ2 T cells after treatment with 107G3B5, as Vγ9Vδ2 T cells cocultured with PC-3 spheroids produced higher levels of IFNγ compared to other solid tumor cells lines (Fig. 3e).

We next used holotomographic microscopy to visualize the impact of 107G3B5 on the efficiency of Vγ9Vδ2 T cells to induce apoptosis in Ishikawa solid tumor cells. Consistent with previous results, we observed a rapid accumulation of apoptotic cells over time under 107G3B5 treatment, whereas no apoptotic feature was distinguishable in the control condition (Fig. 3f, Supplementary Movie 3 and 4). The dead cells fraction was strongly increased upon 107G3B5 treatment (Fig. 3g) and was correlated with higher IFNγ production (Fig. 3h).

### 107G3B5 increases interactions between target cells and Vγ9Vδ2 T cells

To further characterize the mechanisms of action of 107G3B5, we assessed its impact on the interactions between target cells and Vγ9Vδ2 T cells. To this end, we used holotomographic microscopy, a live-imaging technique that enables the close monitoring of both target and effector cells at high resolution ^37^. To obtain relevant findings, we used both a hematological malignancy model; SUP-T1 and a solid cancer model, Ishikawa. Upon 107G3B5 treatment, we observed an increased proportion of Vγ9Vδ2 T cells in contact with a target cell in both cell lines (Fig. 4a). Consistently there was an increase in the target cell area covered by Vγ9Vδ2 T cells (Fig. 4b) and the average minimum distance between Vγ9Vδ2 and target cells. Conversely the movement of Vγ9Vδ2 cells was significantly decreased (Fig. 4c and d). These results strongly suggest that 107G3B5 promotes contacts between Vγ9Vδ2 T cells and target cells and stabilizes their interactions. Interestingly, different behaviors have been reported when T cells encounter antigen on the surface of APC (antigen-presenting cells) such as cessation of movement and the formation of a stable immunological synapse or continuous rapid movement from APC to APC ^38^. Accordingly, molecular interactions between Vγ9Vδ2 T cells and target cells are sufficient to induce the formation of an immunological synapse, but effector functions require the full engagement of the γδ TCR by a potent agonist^39^.Overall, we have provided new insights into the mechanism of Vγ9Vδ2 T cell functionality in different tumor cell lines. 107G3B5 improved the quantity and the quality of interactions between target cells and Vγ9Vδ2 T cells over time, with subsequent increased cytotoxicity.

**Fig. 4.**
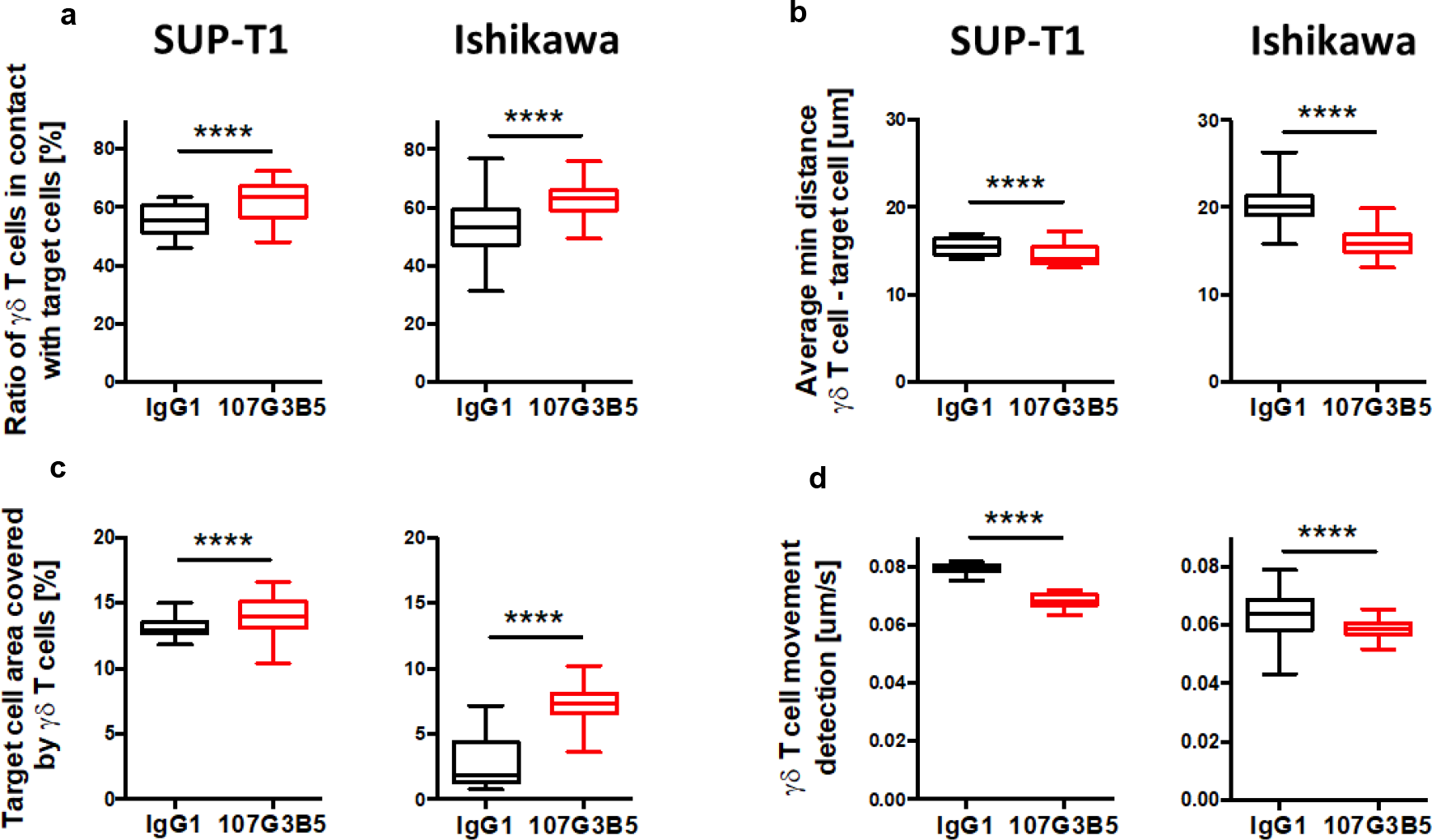
107G3B5 induces a strong increase in Vɣ9Vδ2 TCR binding which is associated with BTN2A1 clustering. (**a**) The ratio of γδ T cells in contact with target cells, (**b**) target cell area covered by γδ T cells in percentage, (**c**) the average minimal distance between γδ T cells centroid and target cells centroid and (**d**) the movement of γδ T cells in um/s were analyzed using a holotomography microscope. These four parameters were analyzed in γδ T cells (Effector: E) coculture with SUPT1 (Target: T; ratio T/E = 1/2) or Ishikawa (ratio 1/3) treated with 10ug/ml of mIgG1 control isotype (in black) or anti-BTN2A1 107G3B5 (in red). Acquisitions were performed each 9min for 20 hours. The boxplots represent for each condition the cumulative results of the acquisition between 4 and 20hours. The statistical significance was established using paired T test. ****p < 0.0001

### 107G3B5 enhances binding of a recombinant Vγ9Vδ2 TCR on HEK293T, SUPT-T1 and Ishikawa cell lines

To gain insight into the underlying molecular mechanisms we assessed whether 107G3B5 can directly affect the binding of a recombinant Vγ9Vδ2 TCR to target cells. First, using HEK293T, SUP-T1 and Ishikawa cell lines, we demonstrated by flow cytometry that 107G3B5 significantly increased Vγ9Vδ2 TCR binding to target cells (Fig. 5a and b). Second, we confirmed this result by confocal microscopy experiments using the SUP-T1 cell line. We observed that the binding of Vγ9Vδ2 TCR did not occur diffusely but in specific areas of the target cells creating clusters of fluorescence (Fig. 5c). Indeed, we observed both mini and microclusters. Using a manual analysis, enabling quantification of the number of miniclusters, we found that 107G3B5 induced a significant increase in the number of Vγ9Vδ2 TCR miniclusters on the target cell surface (Fig. 5d). Consistent with our results, confocal microscopy experiments also showed that the 107G3B5 antibody induced BTN2A1 clustering (Supplementary Fig. 7). Interestingly, treatment of target cells using the potent Vγ9Vδ2 activator HMBPP induced only microclusters of Vγ9Vδ2 TCR on the SUP-T1 surface. Taken together, our results showed that 107G3B5 has its own unique mechanism of action by directly and strongly enhancingVγ9Vδ2 TCR binding to various target cells in a BTN2A-clustering dependent manner.

**Fig. 5.**
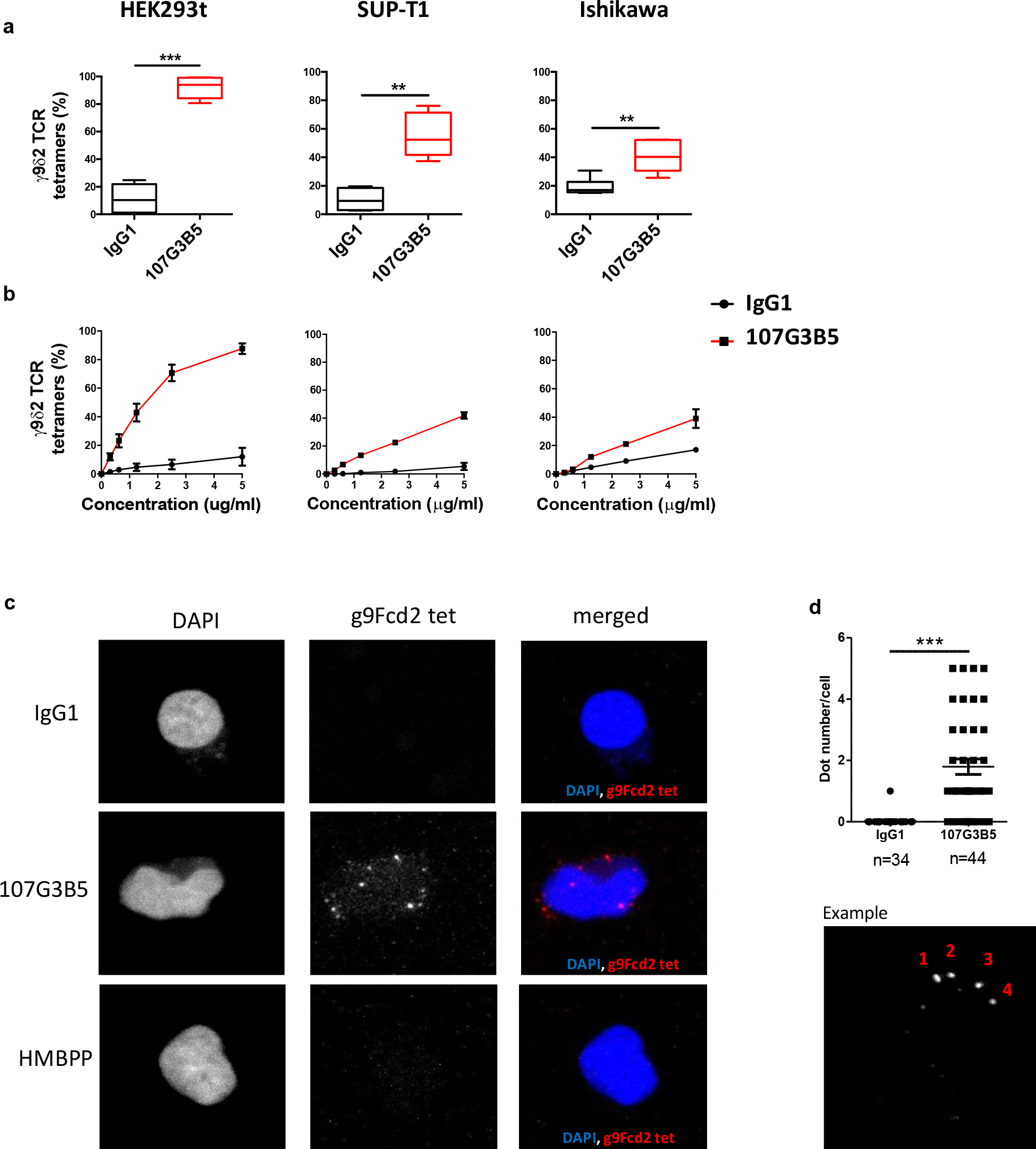
107G3B5 induces a strong increase in Vɣ9Vδ2 TCR binding which is associated with BTN2A1 clustering. (**a,b**) Binding of a recombinant, tetramerized Vγ9Vδ2 TCR to HEK293T IRR, SUP-T1 or Ishikawa, following saturation with 107G3B5 or an isotype control. Binding of Vγ9Vδ2 TCR (5µg/mL) (**a**) or dose dependent binding was monitored by flow cytometry (**b**). (**c**) Confocal microscopy of tetramerized Vγ9Vδ2 TCR binding to SUP-T1, with or without previous saturation with an isotype control,107G3B5 or HMBPP. Evaluation of Vγ9Vδ2TCR binding by quantification of dots observed after treatment with IgG1 control isotype or 107G3B5. Statistical significance was established using Wilcoxon test (**a**) or Mann Whitney test (**d**). **p < 0.01; ***p < 0.001.

To better understand the molecular mechanism, we investigated the effect of 107G3B5 on BTN3A1/BTN2A1 interaction. To this end, HEK293T cells were transfected with BTN3A1-GFP and BTN2A1-mCherry constructs and incubated with agonist 107G3B5, antagonist 7.48 mAbs or control isotype. 107G3B5 was able to significantly increase the colocalization of both molecules, as indicated in yellow in merged images (Supplementary Fig 6a). Indeed, the fraction of total BTN2A1 fluorescence that colocalized with BTN3A1 fluorescence, as indicated by the Manders’ Colocalization Coefficient, was significantly increased on the HEK293T cell line after exposure to 107G3B5 (Supplementary Fig 6b). However, we observed no significant effect of the antagonist antibody 7.48 on BTN2A1/BTN3A1 clustering (Supplementary Fig 6b). To validate this idea, we used BTN2A1KO/BTN3A1KO HEK293T cells transfected with CFP/YFP BTN2A1 and BTN3A1 fusion proteins and incubated with increasing concentrations of 107G3B5 mAb or control isotype. We observed an increasing FRET signal under anti BTN2A-treatment (Supplementary Fig 6c). The interaction of these two molecules is part of the mandatory first steps of Vγ9Vδ2 T cell activation^11^ and our results demonstrate that 107G3B5 promotes the colocalization of both molecules.

### Vγ9Vδ2 T cell activation by 107G3B5 triggers tumor cells pyroptosis

Activation of caspase 3/7 can lead to apoptosis, a non-inflammatory form of programmed cell death or pyroptosis, a form of pro-inflammatory cell death that has been discovered in recent years^40^. Interestingly, pyroptosis is preferentially triggered when CAR T cells have a high affinity for target cells^28^. Because 107G3B5 increases interactions between target cells and Vγ9Vδ2 T cells we wanted to determine whether pyroptosis may contributes to 107G3B5 activity. Pyroptosis is orchestrated by the cleavage of Gasdermin (GSDM) family members such as GSDMD and GSDME. Indeed, GSDME, a 55kDa protein, can be cleaved by caspase-3/7 to produce a 35kDa active form that oligomerizes into the plasma membrane causing pores formation and subsequent cellular pyroptosis^28^. When we co-incubated Vγ9Vδ2 T cells with SUPT1, 697 or Ishikawa cell lines, we observed that 107G3B5 not only increased caspase 3/7 activation, but also increased tumor cell permeabilization via Live/Dead+ marker (Fig. 6a and b). To better understand whether pyroptosis might be involved in this permeabilization, we thought to visualize the GSDME forms by western blot. We observed that the full-length form of GSDME was expressed in the 697 and ishikawa cell lines but was absent in Vγ9Vδ2 T cells. In target/effector co-cultures with control isotype, we observed the full-length form of GSDME but also weakly the N-terminal active form, indicating that Vγ9Vδ2 T cells alone can induce GSDME cleavage. Moreover, the relative expression of the full-length form of GSDME is reduced under 107G3B5 treatment whereas the active form is significantly increased (Fig.6 c and d). Taken together, these results suggest that 107G3B5 mediates the activation of Vγ9Vδ2 T cells, which can mobilize GSDME to induce pyroptosis within the target cell.

**Figure 6.**
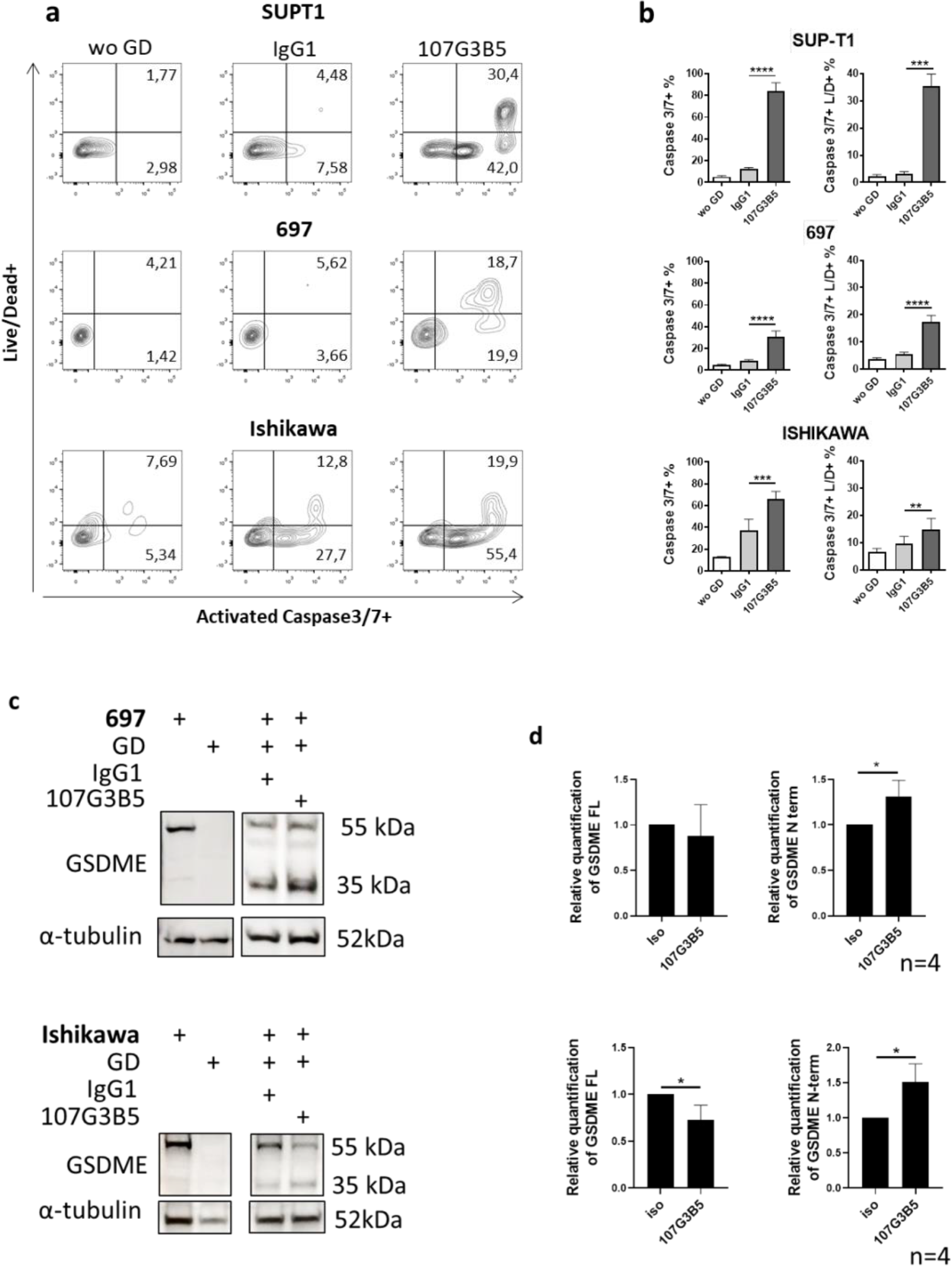
Activation of Vγ9Vδ2 T cells by 107G3B5 induces cancer cell pyroptosis. Vγ9Vδ2 T cells were cocultured with SUPT1, 697 or Ishikawa cells at an effector/target ratio (E/T), 2/1, 2/1 and 3/1, respectively. Cells were treated with 10ug/ml of mIgG1 control isotype or anti-BTN2A1 107G3B5 for 5 hours. The percentage of activated caspase 3/7+ and or permeabilized Live/Dead + tumor cells was determined by flow cytometry. (**a**) Representative staining. (**b**) Data are means ±SD of 3 independent experiments (n>5). The statistical significance was established using a paired T-test. (**c,d**) GSDME and tubuline were analyzed by Western blot. (**c**) Visualization of representative Western blot. (**d**) Data are means of relative quantification ±SD of 4 independent experiments. Statistical significance was established using a paired T-test. \**p*< 0.05; **p < 0.01; ***p < 0.001; ****p < 0.0001.

## Discussion

VγV9δ2 T cells provide a natural bridge between innate and adaptive immunity, responding rapidly and strongly to cancer cells or pathogenic infections. Expression of BTN2A1 is required to trigger activation of Vγ9Vδ2 T cells by pAgs. However, current data on BTN2A1 expression are very sparse. Indeed, BTN2A1 expression has been shown on circulating B, T, NK cells, monocytes and VγV9δ2 T cells from healthy volunteers^13^. Expression on tumor was known for only a few cell lines ^10^. To develop new therapeutic tools targeting BTN2A1, it is necessary to study BTN2A1 in a patient cohort. Therefore, we carefully investigated the expression of BTN2A in normal and tumor tissues. Our results demonstrate that BTN2A1 is ubiquitously expressed at the transcriptomic and protein level. Moreover, BTN2A1 is found on the cell surface of tumor from different origins including primary ALL tumor cells. These data confirm that BTN2A1 can be targeted in a wide range of tumors.

Indeed our results show that in hematological cell lines the activity profile of 107G3B5 was promising in both lymphoid and myeloid malignancies. Next, we confirmed that 107G3B5 enhanced effector functions of Vγ9Vδ2 T cells against primary ALL cells at diagnosis. These findings suggest that BTN2A1 should be considered as a novel immune-checkpoint to target in cancer immunotherapy.

The tumor models used in this study showed different profiles of susceptibility to Vγ9Vδ2 T cell killing, either in basal condition or after stimulation with 107G3B5. In solid tumors, we found that PC-3, a prostate cancer cell line was particularly sensitive to the cytolytic activity of Vγ9Vδ2 T cells after the use of 107G3B5. Therefore, targeting BTN2A1 may be a good alternative to overcome the limited cytotoxic effects of Vγ9Vδ2 T cells on PC-3 cells under zoledronate treatment ^41^.

BTN2A1 is a surface protein, that has been described as a key molecule that binds directly to the γ9 chain of the TCR. However, the molecular mechanisms involved in the regulation of Vγ9Vδ2 T cell activation are still unclear. We have recently shown that BTN2A1 mAbs can inhibit the ability of Vγ9Vδ2 T cells to kill cancer cells^10^. Indeed, BTN2A1 mAbs are able to completely prevent the binding of BTN2A1 to the Vγ9Vδ2 TCR. Surprisingly, we report here for the first time that a unique anti-BTN2A 107G3B5 mAb can induce Vγ9Vδ2 T cell activation. Our work demonstrates that the binding of 107G3B5 to BTN2A1 expressing cells increases the interaction of recombinant Vγ9Vδ2 TCR. To better understand the underlying molecular mechanisms of action, we analyzed the impact of 107G3B5 on BTN2A1 clustering and on its association with BTN3A1. We observed that under 107G3B5 treatment, BTN2A1 appears to form clusters at the surface of cancer cells. Given that TCR avidity depends on both the affinity of a single TCR molecule for its ligand and the number of TCR-ligand engagements, we hypothesis that BTN2A1 clusters increase the avidity for Vγ9Vδ2 TCR. We also observed that 107G3B5 treatment increased the interaction between BTN2A1 and BTN3A1. We do not exclude that other associated molecules may be directly required. Our findings support a model in which BTN2A1 is more than a passive interacting molecule, BTN2A1 appears to be an on/off switch molecule for Vγ9Vδ2 T cells. Therefore, BTN2A1 represents a fundamentally different class of immune recognition compared to the well-known antigen presenting molecules such as MHC and MHC-like molecules.

The pAgs bind to the B30.2 intracellular domain of BTN3A1 and this has recently been defined as a key step in initiating an intracellular heterodimeric association between BTN3A1 and BTN2A1^11^. We demonstrate here that 107G3B5 is also able to enhance BTN2A1 and BTN3A1 association. However, we did not ruled out the possibility that their intracellular domains is involved in such phenomena. The involvement of intracellular or extracellular domains in the observed colocalization induced by the 107G3B5 should be further investigated.

Standards in onco-immunology for the visualization and characterization of lymphocyte’s effector functions and cytotoxic effects on cancer cells are based on the immunolabeling of specific membrane or intracellular proteins with fluorescent antibodies or on the use of chemical dyes. This has two disadvantages: first, the labels alter cellular functions^42, 43^; second, the excitation light used to stimulate fluorescence produces phototoxicity and photobleaching^43^. Holotomographic microscopy (HTM) used in this study does not have these drawbacks as it is label-free and uses extremely low power light^37^. HTM reports the refractive index distribution of the observed sample and enables previously impossible long-term time lapse imaging of lymphocytes and tumor cells. Combined with advanced computer vision, HTM provides unique quantitative information, particularly in this study, on the dynamic interactions between effector and target cells. Specifically, we are able to show that 107G3B5 promoted a long-lasting anti-tumor response of Vγ9Vδ2T cells and illustrate the effect of 107G3B5 on the establishment and stabilization of the effector-target cell interaction. One of the promising applications of HTM is to monitor, quantify and compare the effects of multiple immunotherapeutic treatments on malignant cell death over time.

Despite the growing importance of immunotherapy in cancer, its applications is severely limited by the fact that a high percentage of patients do not respond to these treatments or respond only transiently. Many cancer tumors are not inflamed or “cold,” meaning that the immune system does not recognize them. Induction of apoptosis has long been considered as one of the most important strategy for cancer treatment, as the regulation of apoptosis allows tumors elimination. In the last decade, many new forms of non-apoptotic cell death including pyroptosis has been discovered. This offers new strategies for cancer treatment. Interestingly, the immunogenicity of tumor cells can be maximized by inducing non-apoptotic cell death. Indeed, recent studies have found that GSDME-mediated pyroptosis enhances the phagocytic function of tumor associated macrophages on tumor cells and increases the number and function of tumor-infiltrating T CD8+ /NK cells as well^27^. In melanoma patients, gene signature associated to the increased pyroptosis was correlated with a better prognosis and was predictive of response to anti-PD-1 immunotherapy. Consequently, the understanding of pyroptosis could greatly enhance the efficacy of immunotherapy.

Cytotoxic cells such as Natural Killer (NK) cells, CD8+ T cells and CAR T cells induce tumor cells pyroptosis through pyroptosis^27, 28^. Importantly here we show here for the first time that Vγ9Vδ2 T cells are able to convert non-inflammatory apoptosis into pyroptosis in GSDME-expressing cancer cell lines. Interestingly, pyroptosis induced by the GSDME cleavage is strongly enhanced by anti-BTN2A1 treatment.

Tumor cell death by pyroptosis is often dampened by epigenetic silencing of GSDME due to hypermethylation of the GSDME promoter which is found in approximately 52-65% of primary cancers ^44^. However, treatment with the demethylating agent 5-aza-2’-deoxycytidine or decitabine can induce the expression of GSDME ^45, 46^. γδ T cells are attractive effector cells for cancer immunotherapy because they exert potent cytotoxicity against a wide range of cancer cells. Although these approaches have been well tolerated, the clinical responses have generally been infrequent and of short duration. In this context, enhancing immunogenic cell death such as pyroptosis appears to be a promising strategy to improve the efficacy of γδ T cell-based immunotherapy.

In conclusion, our results demonstrate that targeting BTN2A is a promising option as it (i) allows the sensitization of ALL in both allogeneic and autologous settings as well as solid tumor models to Vγ9Vδ2 T cells, (ii) by quantitatively and qualitatively improving the interaction between target and effector cells, associated with (iii) BTN2A1 clustering and BTN3A1 colocalization, triggering (iv) the conversion of cancer cell apoptosis into inflammatory pyroptosis. Future work is needed to determine the potential role of other immune cytotoxic cell subsets such as CD8 T cells, B cells and NK cells that express the target. Finally, our findings are the first to demonstrate the great potential of targeting BTN2A for the treatment of cancer. Additional preclinical data will guide the future development of anti BTN2A treatments. It is tempting to speculate that this discovery may help to improve clinical outcomes for patients with cancers, including non-inflammatory tumors that have so far failed to respond to the most promising immunotherapy treatments.

## Material and methods

### Patients

PBMCs from 28 adult ALL patients at diagnosis (10 B-ALL, 9 T-ALL and 9 Ph+ ALL) were analyzed for their BTN expression level, and some were also tested in functional assays (7 B-ALL, 7 T-ALL and 3 Ph+ ALL). Heparinized blood from ALL patients was obtained from the Hematology Department of the Institut Paoli Calmettes. Informed consent was obtained from all donors in accordance with the Declaration of Helsinki, and the study was approved by our Institutional Review Board (BTN-LAL-IPC2021-049 - Immunomodulation dans les leucémies aigues - June 22, 2021). Baseline characteristics of ALL patients are provided in supplementary Table 1.

### Cell lines

Cell lines derived from acute lymphoblastic leukemia (697, RS4;11, NALM-6, HPB-ALL, SUP-T1; CCRF-CEM), acute myeloid leukemia (GF-D8, HL-60, Kasumi-3, MOLM-13, SKM-1), anaplastic large-cell lymphoma (COST, KARPAS-299, SUDHL-1), non-Hodgkin lymphoma (DAUDI), CML (K562), breast cancer (MCF-7), colorectal cancer (HT-29), prostate cancer (PC-3), pancreatic cancer (PANC-1), cervical cancer (DoTc2, CaSki), and endometrial cancer (Ishikawa) were obtained from the American Type Culture Collection or from DSMZ. In addition, we used HEK293T for microscopy experiments and binding assays.

697, GF-D8, Kasumi-3, SKM-1 were cultured in RPMI 1640 with 20% Fetal Bovine Serum (FBS). PC-3, PANC-1, DoTc2, CaSki and Ishikawa were cultured in DMEM 20% FBS. RS4;11, NALM-6, HPB-ALL, SUP-T1; CCRF-CEM, HL-60, MOLM-13, COST, KARPAS-299, SUDHL-1, DAUDI, K562, MCF-7, HT-29 were cultured in RPMI 1640 with 10% FBS.

Generation of BTN3AKO/BTN2AKO HEK293T cells using CRISPR-Cas9-mediated inactivation of BTN2A and/or BTN3A as previously described^10^. The absence of BTN2A and/or BTN3A was confirmed by flow cytometry.

All cell lines were tested negative for mycoplasma.

### Tumor spheroids generation

MCF-7, HT-29, DoTc2, PC-3, CaSki and Ishikawa cell lines were cultured to form spheroids. Depending on the type of functional assay, 1×10^4^ to 2 x10^5^ cells were plated in 96 well low-attachment U-bottom plates (Costar) in cell culture medium containing 10% FBS. Cells were incubated for 5 days at 37°C and 5% C0_2_ before use in functional assays.

### Expansion of Vγ9Vδ2 T cells

PBMCs from healthy volunteers (HV) were provided by the local Blood Bank (French Blood Establishment). PBMCs from ALL patients and HV were isolated by density gradient (Lymphoprep, REF). For ALL patients, cells were frozen until use. For establishment of allogenic Vγ9Vδ2 T cells, fresh PBMCs from HV were stimulated with zoledronate 1µM (Sigma-Aldricht), and rhIl-2 (Miltenyi) at day 0. From day 5, rhIL-2 was renewed every two days and cells were kept at 1.5×106 /ml for 14 days. Only cell cultures that reached more than 80% of Vγ9Vδ2 T cells purity at day 14 were selected and frozen until use. Allogeneic Vγ9Vδ2T-cells were thawed and cultured overnight in rhIL-2 (200 UI/mL) for use in functional assays.

To generate autologous Vγ9Vδ2 T cells, frozen PBMCs from ALL patients were stimulated by adding rhIL-15 (10ng/mL) using the same protocol, as previously described to enhance proliferative capacities of Vγ9Vδ2 T cells^47, 48^, including in PBMCs from renal cell carcinoma patients^17^ or from AML patients ^49^. Fresh autologous expanded cells were then used for functional assays, depending on the quantity of Vγ9Vδ2 T cells. The fold increase of viable Vɣ9Vδ2 T cells was calculated according to the formula: (%Vɣ9Vδ2 at day 14 × total cell number at day 14)/ (% Vγ9Vδ2 at day 0 × total cell number at day 0).

### Transcriptomic expression of BTN2A and BTN3A1

RNA-seq of human normal or tumor tissue samples were respectively selected from the Genotype-Tissue Expression (GTEx) Project portal, and from the PanCancer Genome Atlas, respectively.

For scRNA-Seq analysis multiple datasets of normal and tumor origin were analyzed (bone marrow from healthy volunteers and AML patients: GSE1166256-6; colorectal cancer patients: GSE144735; breast normal and tumor tissues: GSE164898; non squamous cells Lung carcinoma Investigation description E-MTAB-8107; hepatocellular carcinoma GSE 140228; basocellular carcinoma GSE11625; renal Cell Carcinoma phs002065.v1.p1).

For bulk RNA-seq datasets, data were log2 normalized, quantile normalized, and collapsed by using median values, using Phantasus **v1.11.0** software.

For sc-RNA seq, data were log2, batch effect was corrected using mutual nearest neighbors (MNN)method, only transcripts found in more than 5 cells and expressed in more than 200 cells were analyzed. The fraction of transcripts from mitochondrial genes was set up to 5% for normal tissues study and 10% for the tumor tissue study. Identification of subpopulations was realized using UMAP algorithm and was analyzed using Bioturing software.

### Generation of anti-human BTN2A and BTN3A mAbs

To generate anti-BTN3A 20.1 and anti-BTN3A 108.5 mAbs, BALB/c mice were immunized with a soluble BT3.1-Ig fusion protein, as previously described^50^.

Anti-BTN2A mAbs were produced by mouse immunization using a recombinant human BTN2A1-Fc fusion protein (R&D Biotech), as described in patent WO2019057933A1 (clone 7.48) and patent WO2020188086A1.

### ELISA

Vγ9Vδ2 T cells and target cells were cocultured with at an Effector:Target (E:T) ratio of 10:1. Supernatants were collected and frozen at −80°C for further cytokine measurement. Supernatants were further analyzed for TNFα and/or IFNγ production using respectively BD OptEIA Human TNFα and IFNγ ELISA kits (BD Biosciences). Flat-bottom 96-well plates (COSTAR) were coated with capture Ab overnight andthen blocked with PBS 10% FCS for 1h. After 2h of incubation with standards or supernatants, plates were incubated with detection Ab and streptavidin HRP and then developed using 3,3’,5,5’-tetramethylbenzidine (TMB) Liquid Substrate System (BD biosciences). The reaction was stopped withHCL 2N, and the absorbance was quantified within 30 minutes using a CLARIOstar microplate reader at a wavelength of 450 nm.

### Flow cytometry

Relative surface expression of BTN2A (107G3B5 and 7.48 mAbs) and BTN3A (20.1 and 108.5 mAbs) on cancer cell lines and ALL blasts was assessed by flow cytometry using calibrator beads allowing quantification of antigen density (Quantum™ Simply Cellular^®^ (QSC) kits, Bang Laboratories). To avoid bias related to modulation of BTN surface expression in correlation analysis, relative quantification of BTN was performed concurrently withwith functional assays.

For analysis of CD107 expression, Vγ9Vδ2 T cells and target cells were cocultured in a Effector:Target (E:T) ratio of 1:1 with anti-CD107a, anti-CD107b and Golgi stop, with 107G3B5 or isotype control (1 µg/mL). After 4 hours, cells were harvestedand analyzed by flow cytometry. To study IFNγ and TNFα production, cells were permeabilized with Permwash (BD Biosciences) to allow intracellular staining. Cells were acquired on a FACS CANTO II and analysed using Flowjo software.

### Apoptosis assays

#### Cell lines and primary ALL blasts

Target cells were first labelled with the Cell Proliferation Dye eFluor™ 670 (LifeTechnologies) and then cocultured for4, 6 or 10 hours with effector cells at an E:T ratio of 5:1 (cell lines), 10:1 or 15:1 (primary ALL blasts), in the presence of an isotype control, 107G3B5 or anti-BTN3A 20.1 agonist mAb (1 or 10 µg/mL). In caspase assays with cancer cell lines, target cells preincubated overnight (O/N) with zoledronate (45 µM) were used as a positive control. At the end of coculture, target cells were labeled with CellEvent™ caspase-3/7 green detection reagent for 30 min and analyzed by flow cytometry (FACSCanto II). The percentage of specific apoptosis was calculated using the formula [(experimental – spontaneous caspase 3/7 activation)/ (100 – spontaneous caspase 3/7 activation) × 100].

#### Tumor spheroids

Ishikawa, HT-29, PC-3 and CaSki were cocultured with Vγ9Vδ2 T cells from 6 HV for 10 hours at an E:T ratio of 5:1, in the presence of an isotype control or 107G3B5 (10µg/mL). Caspase-Glo 3/7 reagent was added at the end of the coculture and was incubated for 1 hour. Luminescence was quantified within 30 minutes using a CLARIOstar microplate reader.

### 3-dimensional (3D) tumor spheroids live-cell imaging

After 5 days of spheroid generation, PC-3 were incubated for 4 hours in serum-free medium containing calcein (Sigma) at a final concentration of 2 µM. PC-3 spheroids were then washed extensively and cocultured with Vγ9Vδ2 T cells from 6 HV for 48 hours at an E:T ratio of 5:1, in presence of an isotype control or 107G3B5 (10µg/mL).

Real-time calcein intensity was monitored using fluorescence live-cell imaging combined with a phase-contrast setup using the 4×objective (Sartorius AG, Gottingen, Germany). Images were acquired hourly and analyzed using Incucyte software and ImageJ.

### Live-cell imaging of effector and target cell interactions

The target cells (T); HEK293T, SUP-T1 and Ishikawa; were cultured O/N or 1h for SUP-T1 on dishes (Ibidi, Cat.No: 80136) precoated with poly-L-lysine. Non-adherent cells were removed by washing with PBS. Vγ9Vδ2 T cells expanded from healthy volunteers were added to the target cells (E:T ratio = 2:1 for HEK293T and SUPT-1, 3:1 for Ishikawa) in 1 ml of RMPI 10% SVF. Control isotype or 107G3B5 was added to the final concentration of 10 µg/mL. Dishes were incubated at 37°C with 5% CO_2_ within Tokaihit stage top incubator throughout the acquisitions, which were performed with a CX-A automated microscope. The CX-A (Nanolive) consists of an holotomographic microscope coupled to an automated stage that allows grid scanning and multi-wel acquisition. The microscope uses a 60X air objective (NA = 0.8) at a wavelength of λ = 520 nm (Class 1 low power laser, sample exposure 0.2 mW/mm^2^) and USB 3.0 CMOS Sony IMX174 sensor, single field of view 90 × 90 × 30 μm, axial resolution 400 nm, and maximum temporal resolution of 0.5 3D RI volume per second. The theoretical sensitivity of the RI measurement was 2.71 × 10^−4^. The time lapse acquisition regime was one image every 9 minutes for 20 hours and was the same for all imaging experiments. The dynamics of the lethal fraction was realized as previously described ^51^, using ML-based dead cell identification.

### Quantification of effector/target cells interactions

Quantifications were performed using proprietary computer vision developed at Nanolive. This analysis consists of identifying each cell object and its type (effector cells on one side, target cells on the other) and to quantifying the dynamic interactions between the two populations. The dynamic evolution of four parameters was exploited; the ratio of γδ T cells in contact with target cells, the average minimum distance between γδ T cells and target cell centroids, the percentage of target cell area covered by γδ T cells and the global movement speed of γδ T cells in um/sec. The lethal fraction of target cells, i.e. the cumulative number of dead cells over time, was also quantified. To do this, we used machine learning to train the computer to distinguish the typical shape and texture signatures of living and of dead target cells observed by HTM.

### Recombinant Vγ9Vδ2 TCR binding assays

A recombinant Vγ9Vδ2 TCR (clone G115)^52^ with a C-terminal Fc-tag or His-tag on the Vγ9 chain was produced in CHO cells, then purified using protein A column chromatography followed by buffer-exchange into PBS. Vγ9Vδ2 TCR were biotinylated using protein labeling kit according to the manufacturer’s instructions (Thermofisher). Biotinylated proteins were then tetramerized with streptavidin-APC (Bio-legend) at 4:1 molar ratio by adding the streptavidin over 10 times intervals to maximize tetramer formation. HEK293T, SUP-T1 or Ishikawa cells were saturated with 107G3B5 or isotype control (10 µg/mL) for 4 hours at 37°C. The cells were then stained with recombinant tetrameric Vγ9Vδ2 TCR-Fc at 10 µg/ml for 30min at 37°C. After two washes, AF647 mean fluorescence intensity (MFI) was assessed by flow cytometry (Fortessa).

### Confocal microscopy

To study the colocalization of BTN2A1 and BTN3A1, HEK293T cells were transiently transfected with either with BTN3A1 GFP or BTN2A1 mcherry using PEI max reagent (Fisher Scientific), according to the manufacturer’s instructions. For both constructs, a linker was inserted between the BTN2A1/BTN3A1 C-terminal end and the -eGFP or -mCherry molecules in order to provide flexibility and not interfere with molecular function. The next day, the cells were plated on precoated coverslips with poly-L-lysine. After overnight incubation, cells were preincubated with either an isotype control or anti-BTN2A 107G3B5/7.48 or anti-BTN3A 20.1 mAb (10µg/mL) during 1 hour. After two washes, cells were fixed with 3% paraformaldehyde in phosphate-buffered saline (PBS pH 7.4), saturated with 2% BSA in phosphate-buffered saline (PBS pH 7.4) for 2 hours and labeled with DAPI for 2min. After 2 washes with phosphate-buffered saline (PBS pH 7.4) and 1 wash with deionized water, coverslips were mounted with prolong Gold (Invitrogen). Cells were imaged using an inverted ZEISS LSM 880-Spectral-Airyscan with the 63X objective. BTN2A1 and BTN3A1 distribution and colocalization were analyzed using ImageJ software and the JACoP plugin.

In order to visualize and quantify the colocalization of BTN2A and BTN3A and the binding of the Vγ9Vδ2 TCR on a more relevant model, SUP-T1 cells were preincubated with either an isotype control or 107G3B5 (10µg/mL) or HMBPP (10µM) for 4 hours at 37 °C. After two washes, cells were incubated with either an anti BTN2A 7.48 AF647 mAb or anti-BTN3A 103.2 AF488 mAb (10µg/mL) for 1 hour at 4°C or with Vγ9Vδ2 TCR-Fc at 10 µg/ml for 30min at 37°C. After two washes, cells were fixed, then mounted with prolong Gold and covered with a coverslip overnight. Cells were imaged using an inverted ZEISS LSM 880-Spectral-Airyscan with the 63X objective.

### Western blotting

The cells were lysed in M2 lysis buffer. The protein concentrations were determined using Bradford reagent. Then, the proteins were run on an SDS-PAGE gel and transferred to PVDF membrane which were blocked in 5% milk and probed with antibodies overnight with antibodies: anti α-tubulin and anti GSDME (cell signaling). Secondary antibody conjugated with biotin and third antibody conjugated with horseradish peroxidase were followed by enhanced chemiluminescence (Cytivia). Results were confirmed by at least three independent experiments and quantified using Fiji.

### Statistics

All analyses were performed using the GraphPad Prism 5.0 software package.

The normality distribution of the data was determined by using the D’Agostino test. Appropriate statistical tests were used as indicated in the figure legends. All replicates’ data are biological except for ELISA experiments, where technical replicates are indicated.

## Acknowledgments

We thank the IPC Immunomonitoring platform and the the IPC Tumor Bank (authorization number AC-2007-33, granted by the French Ministry of Research). We thank the Timone Microscopy platform PIVMI from the Center for Cardiovascular and Nutrition Research. The team “Immunity and Cancer” was labeled “FRM DEQ20180339209” (for D.O.). A.C.L.F was funded by the Poste d’Accueil INSERM from November 2019 to October 2021. We would like to thank Mauro Modesti for advice.

## Author contributions

A.C.L.F, C.I., C.E.C., M.F. and D.O. conceived and designed the study. A.C.L.F, C.I., N.B., L.G., S.F., F.O., A.L.R, L.C., L.D., C.D, A.A. and A.B. performed experiments. A.C.L.F, C.I., N.B., L.G., L.C., L.D., C.D., C.E.C., M.F. and D.O. interpreted the data. N.V. treated patients and coordinated the study. A.C.L.F, C.I., C.E.C, G.G, M.F. and D.O. coordinated the research and wrote the manuscript. All authors critically reviewed the manuscript.

## References

1. Wilcox, R. A. et al. Ligation of CD137 receptor prevents and reverses established anergy of CD8+cytolytic T lymphocytes in vivo. Blood 103, 177–184 (2004).

2. Gao, Y. et al. γδ T Cells Provide an Early Source of Interferon γ in Tumor Immunity. J. Exp. Med. 198, 433–442 (2003).

3. Tosolini, M. et al. Assessment of tumor-infiltrating TCRVγ9Vδ2 γδ lymphocyte abundance by deconvolution of human cancers microarrays. Oncoimmunology 6, e1284723 (2017).

4. Gentles, A. J. et al. The prognostic landscape of genes and infiltrating immune cells across human cancers. Nat. Med. 21, 938–945 (2015).

5. Gober, H.-J. et al. Human T cell receptor gammadelta cells recognize endogenous mevalonate metabolites in tumor cells. J. Exp. Med. 197, 163–168 (2003).

6. Tanaka, Y. et al. Natural and synthetic non-peptide antigens recognized by human γδ T cells. Nature 375, 155–158 (1995).

7. Mullen, P. J., Yu, R., Longo, J., Archer, M. C. & Penn, L. Z. The interplay between cell signalling and the mevalonate pathway in cancer. Nat. Rev. Cancer 16, 718–731 (2016).

8. Sandstrom, A. et al. The intracellular B30.2 domain of Butyrophilin 3A1 binds phosphoantigens to mediate activation of human Vγ9Vδ2 T cells. Immunity 40, 490–500 (2014).

9. Yang, Y. et al. A Structural Change in Butyrophilin upon Phosphoantigen Binding Underlies Phosphoantigen-Mediated Vγ9Vδ2 T Cell Activation. Immunity 50, 1043–1053.e5 (2019).

10. Cano, C. E. et al. BTN2A1, an immune checkpoint targeting Vγ9Vδ2 T cell cytotoxicity against malignant cells. Cell Rep. 36, 109359 (2021).

11. Hsiao, C.-H. C., Nguyen, K., Jin, Y., Vinogradova, O. & Wiemer, A. J. Ligand-induced interactions between butyrophilin 2A1 and 3A1 internal domains in the HMBPP receptor complex. Cell Chem. Biol. (2022) doi:10.1016/j.chembiol.2022.01.004.

12. Yuan, L. et al. Phosphoantigens are Molecular Glues that Promote Butyrophilin 3A1/2A1 Association Leading to Vγ9Vδ2 T Cell Activation. 2022.01.02.474068 Preprint at https://doi.org/10.1101/2022.01.02.474068 (2022).

13. Rigau, M. et al. Butyrophilin 2A1 is essential for phosphoantigen reactivity by γδ T cells. Science (2020) doi:10.1126/science.aay5516.

14. Karunakaran, M. M. et al. Butyrophilin-2A1 Directly Binds Germline-Encoded Regions of the Vγ9Vδ2 TCR and Is Essential for Phosphoantigen Sensing. Immunity 52, 487–498.e6 (2020).

15. Corvaisier, M. et al. V gamma 9V delta 2 T cell response to colon carcinoma cells. J. Immunol. Baltim. Md 1950 175, 5481–5488 (2005).

16. Cordova, A. et al. Characterization of human γδ T lymphocytes infiltrating primary malignant melanomas. PloS One 7, e49878 (2012).

17. Viey, E. et al. Phosphostim-activated gamma delta T cells kill autologous metastatic renal cell carcinoma. J. Immunol. Baltim. Md 1950 174, 1338–1347 (2005).

18. Kabelitz, D., Wesch, D., Pitters, E. & Zöller, M. Characterization of tumor reactivity of human V gamma 9V delta 2 gamma delta T cells in vitro and in SCID mice in vivo. J. Immunol. Baltim. Md 1950 173, 6767–6776 (2004).

19. Santolaria, T. et al. Repeated systemic administrations of both aminobisphosphonates and human Vγ9Vδ2 T cells efficiently control tumor development in vivo. J. Immunol. Baltim. Md 1950 191, 1993–2000 (2013).

20. Lança, T. et al. The MHC class Ib protein ULBP1 is a nonredundant determinant of leukemia/lymphoma susceptibility to γδ T-cell cytotoxicity. Blood 115, 2407–2411 (2010).

21. Gomes, A. Q. et al. Identification of a panel of ten cell surface protein antigens associated with immunotargeting of leukemias and lymphomas by peripheral blood gammadelta T cells. Haematologica 95, 1397–1404 (2010).

22. Gertner, J., Wiedemann, A., Poupot, M. & Fournié, J.-J. Human gammadelta T lymphocytes strip and kill tumor cells simultaneously. Immunol. Lett. 110, 42–53 (2007).

23. Gertner-Dardenne, J. et al. Human Vγ9Vδ2 T Cells Specifically Recognize and Kill Acute Myeloid Leukemic Blasts. J. Immunol. 188, 4701–4708 (2012).

24. Yazdanifar, M., Barbarito, G., Bertaina, A. & Airoldi, I. γδ T Cells: The Ideal Tool for Cancer Immunotherapy. Cells 9, (2020).

25. Sebestyen, Z., Prinz, I., Déchanet-Merville, J., Silva-Santos, B. & Kuball, J. Translating gammadelta (γδ) T cells and their receptors into cancer cell therapies. Nat. Rev. Drug Discov. 19, 169–184 (2020).

26. Du, T. et al. Pyroptosis, metabolism, and tumor immune microenvironment. Clin. Transl. Med. 11, e492 (2021).

27. Zhang, Z. et al. Gasdermin E suppresses tumour growth by activating anti-tumour immunity. Nature 579, 415–420 (2020).

28. Liu, Y., et al. Gasdermin E-mediated target cell pyroptosis by CAR T cells triggers cytokine release syndrome. Sci. Immunol. 5, eaax7969 (2020).

29. Moreno, H., Archetti, L., Gibbin, E., Grandchamp, A. E. & Fréchin, M. Artificial Intelligence-Powered Automated Holotomographic Microscopy Enables Label-Free Quantitative Biology. Microsc. Today 29, 24–32 (2021).

30. Salucci, S., Battistelli, M., Burattini, S., Sbrana, F. & Falcieri, E. Holotomographic microscopy: A new approach to detect apoptotic cell features. Microsc. Res. Tech. 83, 1464–1470 (2020).

31. Gundermann, S. et al. A comprehensive analysis of primary acute myeloid leukemia identifies biomarkers predicting susceptibility to human allogeneic Vγ9Vδ2 T cells. J. Immunother. Hagerstown Md 1997 37, 321–330 (2014).

32. Aswald, J. M. et al. Flow cytometric assessment of autologous γδ T cells in patients with acute myeloid leukemia: Potential effector cells for immunotherapy? Cytometry B Clin. Cytom. 70B, 379– 390 (2006).

33. Benyamine, A. et al. BTN3A molecules considerably improve Vγ9Vδ2T cells-based immunotherapy in acute myeloid leukemia. Oncoimmunology 5, (2016).

34. Siegers, G. M. et al. Anti-leukemia activity of in vitro-expanded human gamma delta T cells in a xenogeneic Ph+ leukemia model. PloS One 6, e16700 (2011).

35. Duval, M. et al. Potential antileukemic effect of gamma delta T cells in acute lymphoblastic leukemia. Leukemia 9, 863–868 (1995).

36. Lamb Jr, L., et al. Human γδ^+^ T lymphocytes have *in vitro* graft *vs* leukemia activity in the absence of an allogeneic response. Bone Marrow Transplant. 27, 601–606 (2001).

37. Sandoz, P. A., Tremblay, C., van der Goot, F. G. & Frechin, M. Image-based analysis of living mammalian cells using label-free 3D refractive index maps reveals new organelle dynamics and dry mass flux. PLoS Biol. 17, e3000553 (2019).

38. Dustin, M. L. Stop and go traffic to tune T cell responses. Immunity 21, 305–314 (2004).

39. Favier, B. et al. Uncoupling between immunological synapse formation and functional outcome in human gamma delta T lymphocytes. J. Immunol. Baltim. Md 1950 171, 5027–5033 (2003).

40. Tan, Y. et al. Pyroptosis: a new paradigm of cell death for fighting against cancer. J. Exp. Clin. Cancer Res. CR 40, 153 (2021).

41. Nada, M. H., Wang, H., Workalemahu, G., Tanaka, Y. & Morita, C. T. Enhancing adoptive cancer immunotherapy with Vγ2Vδ2 T cells through pulse zoledronate stimulation. J. Immunother. Cancer 5, 9 (2017).

42. Fei, Y. Y. et al. Real-time, label-free analysis reveals novel low-affinity binding to blood group antigens by Helicobacter pylori. Anal. Chem. 83, 6336–6341 (2011).

43. Hoebe, R. A. et al. Controlled light-exposure microscopy reduces photobleaching and phototoxicity in fluorescence live-cell imaging. Nat. Biotechnol. 25, 249–253 (2007).

44. de Beeck, K. O., Schacht, J. & Van Camp, G. Apoptosis in acquired and genetic hearing impairment. Hear. Res. 281, 18–27 (2011).

45. Akino, K. et al. Identification of DFNA5 as a target of epigenetic inactivation in gastric cancer. Cancer Sci. 98, 88–95 (2007).

46. Wang, Y. et al. Chemotherapy drugs induce pyroptosis through caspase-3 cleavage of a gasdermin. Nature 547, 99–103 (2017).

47. García, V. E. et al. IL-15 enhances the response of human gamma delta T cells to nonpeptide [correction of nonpetide] microbial antigens. J. Immunol. Baltim. Md 1950 160, 4322–4329 (1998).

48. Aehnlich, P., Carnaz Simões, A. M., Skadborg, S. K., Holmen Olofsson, G. & thor Straten, P. Expansion With IL-15 Increases Cytotoxicity of Vγ9Vδ2 T Cells and Is Associated With Higher Levels of Cytotoxic Molecules and T-bet. Front. Immunol. 11, 1868 (2020).

49. Van Acker, H. H. et al. Interleukin-15 enhances the proliferation, stimulatory phenotype, and antitumor effector functions of human gamma delta T cells. J. Hematol. Oncol.J Hematol Oncol 9, (2016).

50. Compte, E., Pontarotti, P., Collette, Y., Lopez, M. & Olive, D. Frontline: Characterization of BT3 molecules belonging to the B7 family expressed on immune cells. Eur. J. Immunol. 34, 2089–2099 (2004).

51. Forcina, G. C., Conlon, M., Wells, A., Cao, J. Y. & Dixon, S. J. Systematic Quantification of Population Cell Death Kinetics in Mammalian Cells. Cell Syst. 4, 600–610.e6 (2017).

52. Allison, T. J., Winter, C. C., Fournié, J. J., Bonneville, M. & Garboczi, D. N. Structure of a human gammadelta T-cell antigen receptor. Nature 411, 820–824 (2001).

