## Supplementary Figures for "Targeting BTN2A1 enhances Vγ9Vδ2 T cell effector functions and triggers tumor cells pyroptosis"

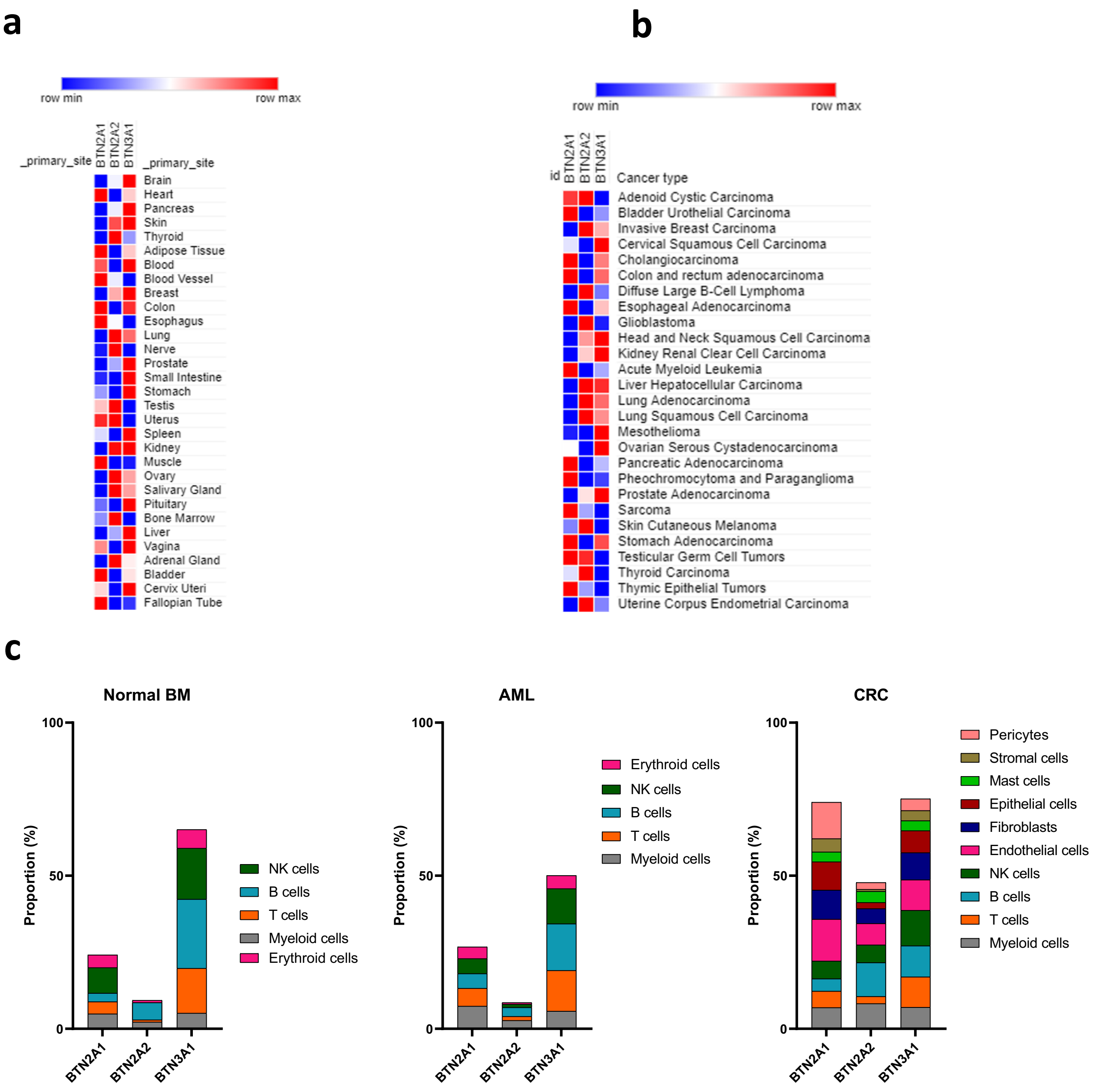


**Supplementary figure 1. BTN2A1 is broadly expressed in normal and tumor cells. (a,b)** BTN2A and BTN3A1 median transcriptomic expression from RNA-seq of 31 human normal tissue samples from the Genotype-Tissue Expression (GTEx) Project **(a)**, and from 27 human tumor tissue samples from the Pancancer genome Atlas **(b)**. **(c)** ScRNA-Seq analysis of BTN2A and BTN3A1 expression in different cell subsets, obtained from public datasets of bone marrow (BM) of Healthy volunteers (HV) or acute myeloid leukemia patients (AML), and from colorectal cancer patients (CRC) . Data are expressed as percentage of positive cells.

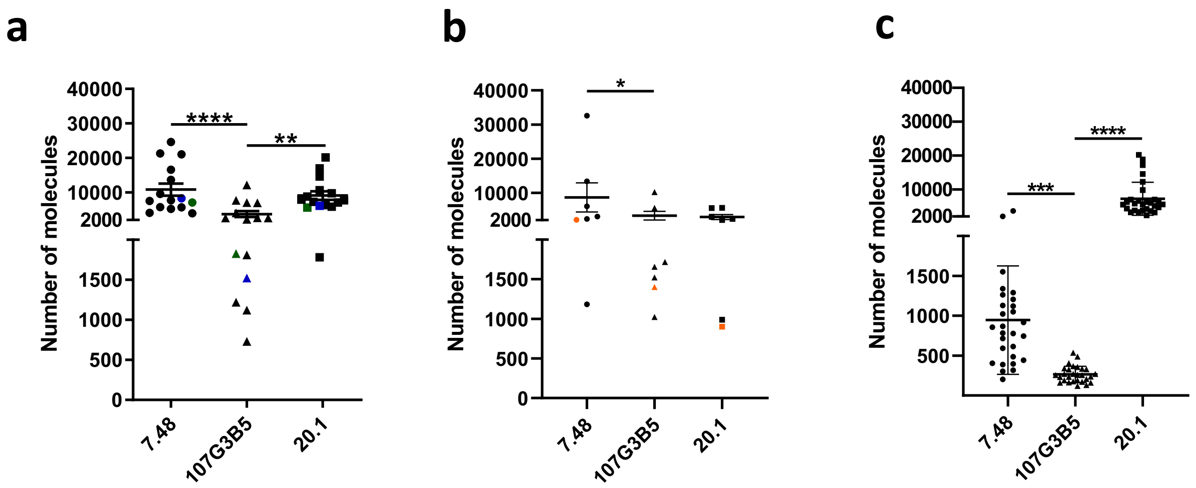


**Supplementary figure 2. BTN2A1 is broadly expressed at the tumor cells surface.** BTN2A and BTN3A surface expression were assessed by flow cytometry on surface of : **(a)** 15 hematological cell lines (697, RS4;11, NALM-6, HPB-ALL, SUP-T1; CCRF-CEM, GF-D8, HL-60, Kasumi-3, MOLM-13, SKM-1, COST, KARPAS-299, SUDHL-1, DAUDI; **(b)** 28 primary ALL blasts (10B-ALL, 9T-ALL, 9Ph+ ALL; **(c)** and 7 solid cancer cell lines (MCF-7, HT-29, DoTc2, PC3, CaSki, PANC-1, ISHIKAWA). 697, SUP-T1 and ISHIKAWA are depicted respectively in green, blue and orange. The relative number of BTN2A and BTN3A molecules were quantified with Molecules of Equivalent Soluble Fluorochrome (MESF) method, using 7.48, 107G3B5 mAbs for BTN2A and 20.1 mAb for BTN3A. The statistical significance was established using using Friedman test with Dunn's post-test **(a,b,c)**. *$p < 0.05$; **$p < 0.01$; ***$p < 0.001$; ****$p < 0.0001$

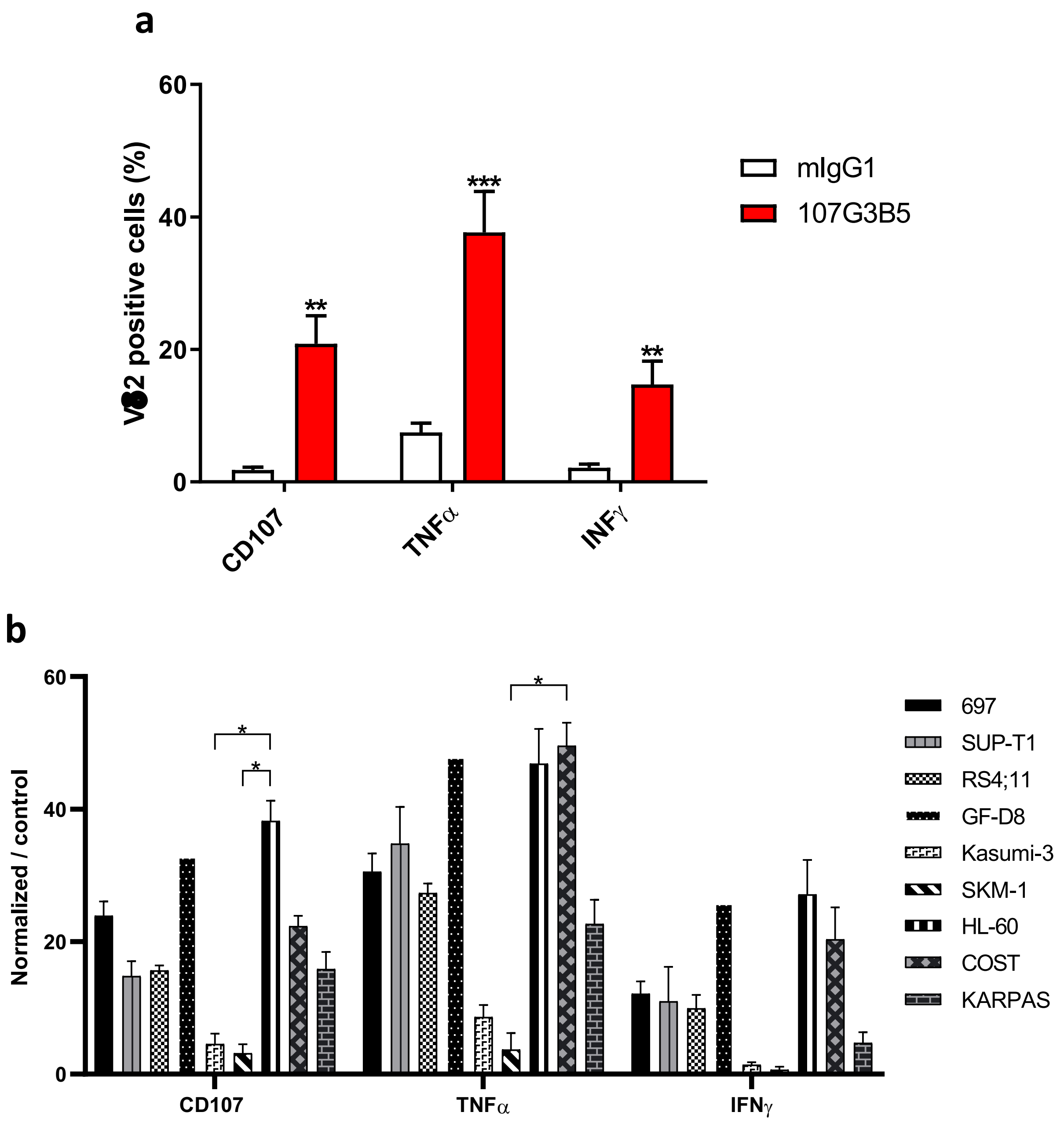


**Supplementary Figure 3. Anti-BTN2A1 107G3B5 increases Vɣ9Vδ2 T cell effectors functions against hematological cell lines. (a,b)** Effector functions of Vγ9Vδ2 T cells (HV, n=3) were assessed after a 4 hours co-culture with 9 hematopoietic cell lines (E:T ratio 1:1) in presence of anti-BTN2A 107G3B5 mAb (10µg/mL) or its isotype control. Bar plots show the percentage of CD107ab+, IFNɣ+ and TNFα+ Vγ9Vδ2 T cells **(a)**, normalized to control condition (**b**, % after stimulation –% in control condition). Mean ± s.e.m. pooled from 2 independent experiments. The statistical significance was established using a paired T-test **(a)**, a Kruskal-Wallis test with Dunn's post-test **(b)** *$p$< 0.05; **$p$ < 0.01; ***$p$ < 0.001

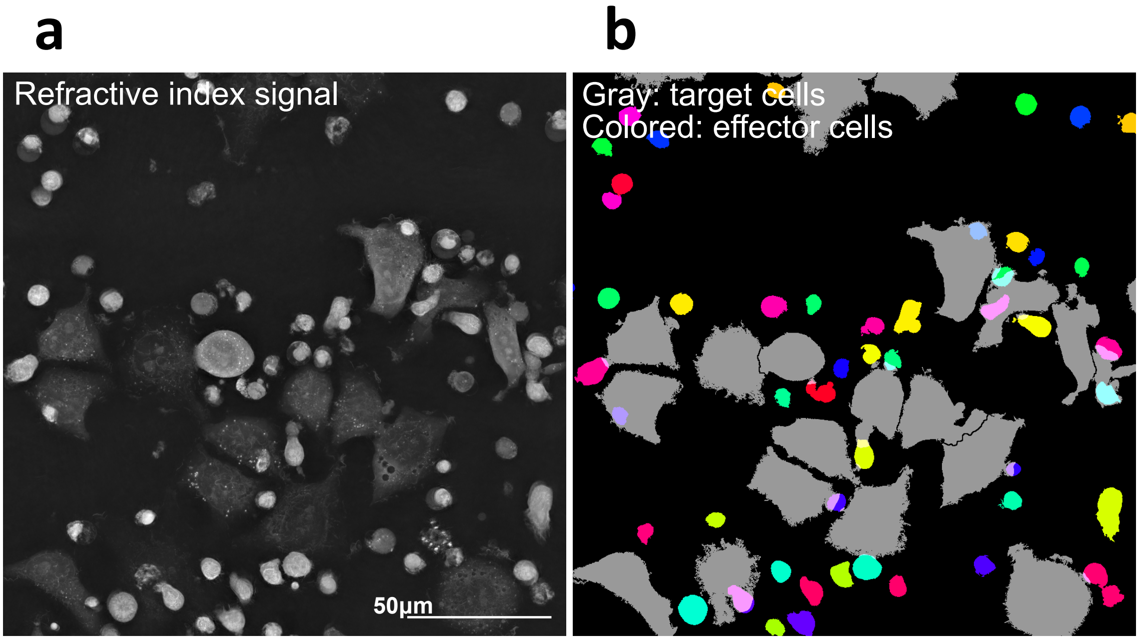


**Supplementary Figure 4. Segmentation of live cells using Refractive Index (RI). (a)** Refractive index signal of γδ T cells (Effector : E) coculture with Ishikawa (Target : T; ratio T/E = 1/3). **(b)** Cell segmentation was computed with EVE Analytics (EA) using the RI signal. Target cells are shown in gray and effector cells are colored.

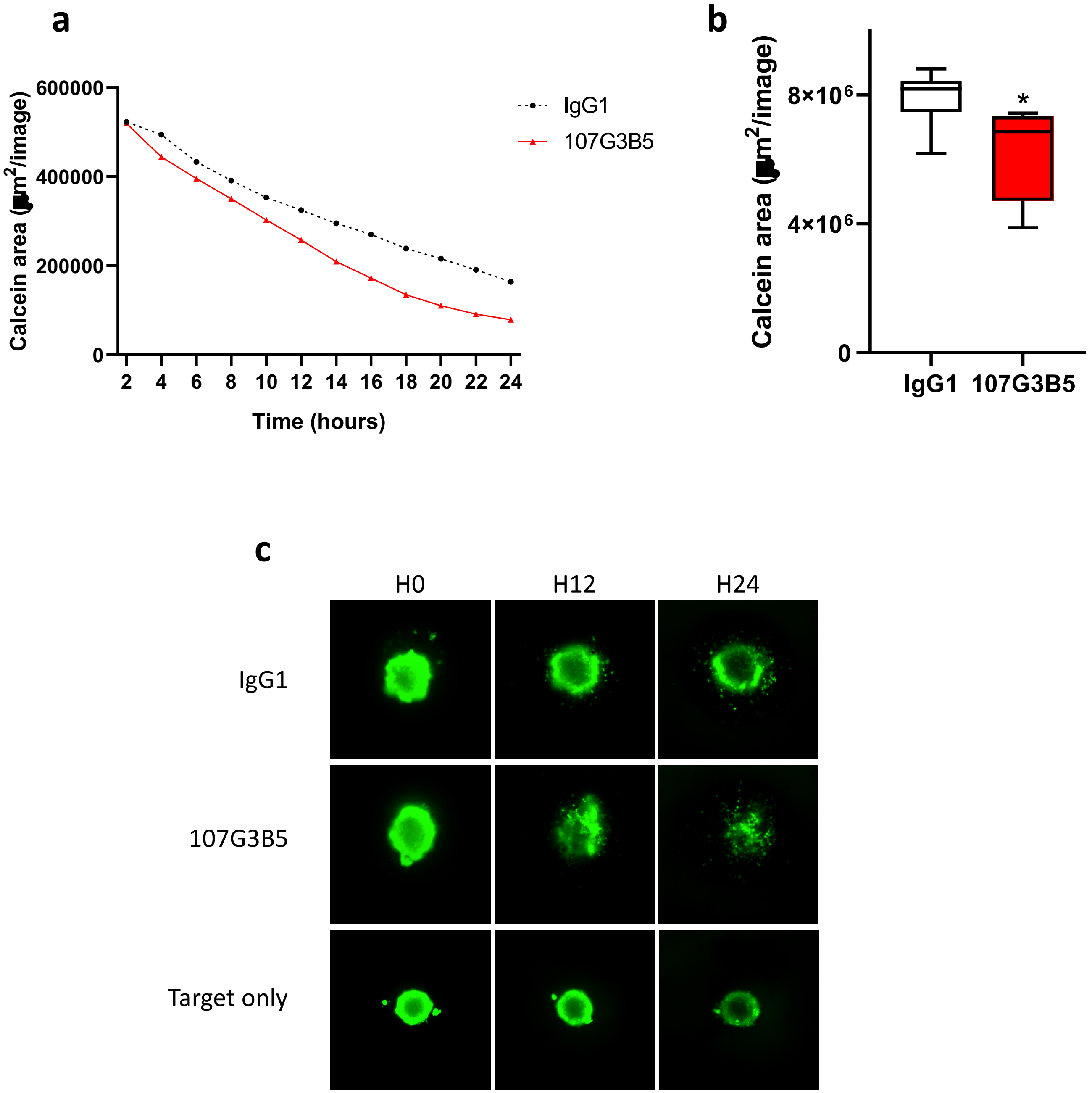


**Supplementary Figure 5. Anti-BTN2A1 sensitizes Vɣ9Vδ2 T cell cytotoxicity against PC3 cell line.** **(a)** Calcein area of spheroid PC3 spheroid, was monitored during a 24-hour coculture with Vγ9Vδ2 T cells from 6 HV (E:T ratio 5:1) in presence of an isotype control anti-BTN3A 20.1 mAb (10µg/mL) or anti-BTN2A 107G3B5 mAb (10µg/mL). **(b)** Box-plot of normalized calcein area after 24-hrs coculture are representative from three independent experiments performed in duplicate. **(c)** Representative scanned live 3D images of PC3 spheroid stained with calcein-AM after 12 and 24 hours of coculture with Vγ9Vδ2 T cells from 1 HV, and after 12 or 24 hours.

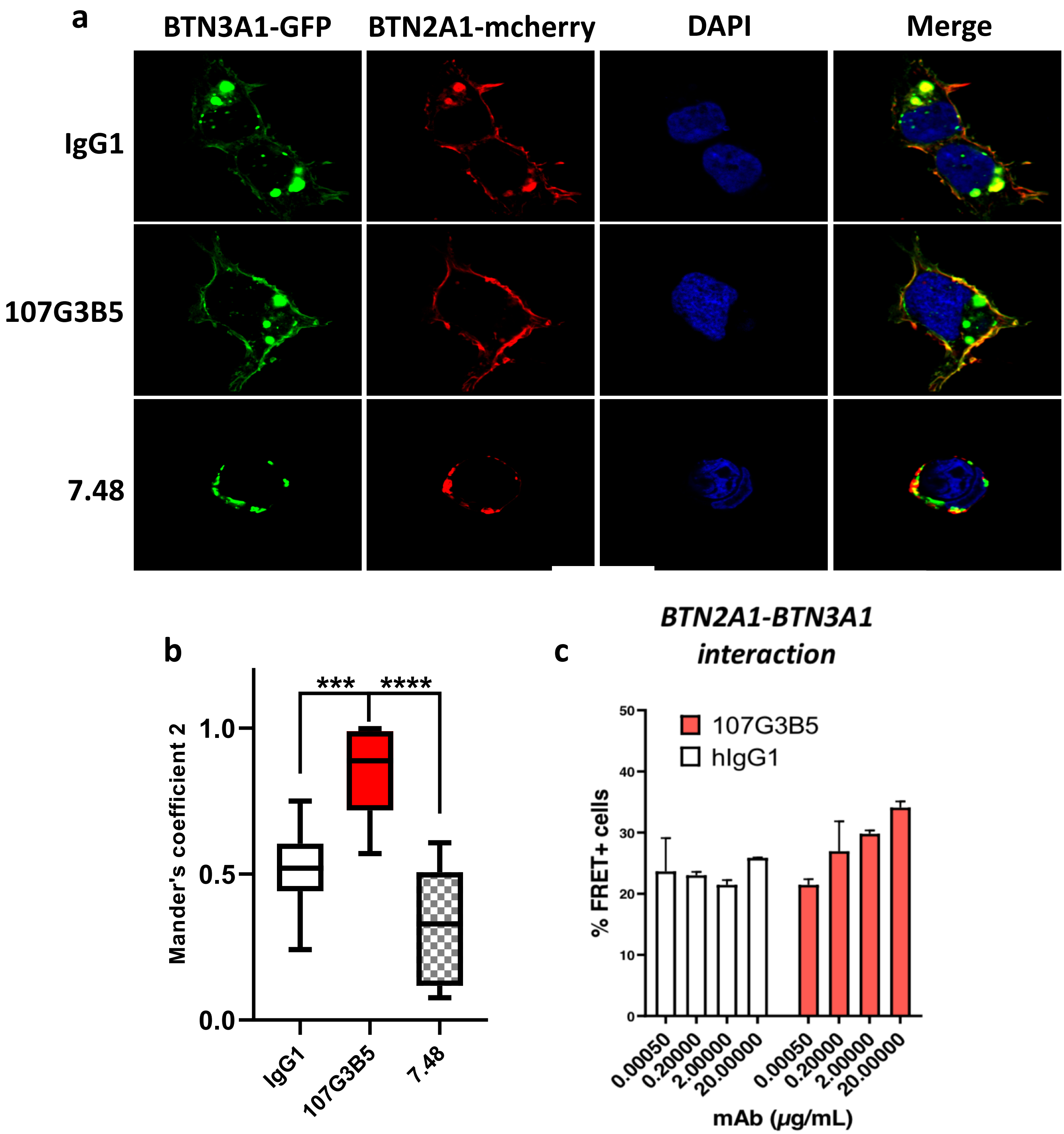


**Supplementary figure 6. Anti-BTN2A1 107G3B5 induces a colocalization of BTN3A1 and BTN2A1.** HEK293T cells were transfected with BTN3A1 GFP and BTN2A1 mCherry and treated with IgG1, 107G3B5 or 7.48 (10ug/ml). The Colocalisation between BTN2A1 and BTN3A1 was visualized by confocal microscopy **(a)** and quantified **(b)** using the Threshold-Manders coefficient (n=17, $*p < 0.05$) obtain with the ImageJ software. **(c)** HEK293T cells were transfected with plasmids encoding BTN2A1-CFP and BTN3A1-YFP, treated overnight with CHX+bafilomycin in presence of 107GB5 or its isotype control. FRET+ cells percentages were assessed by cytometry

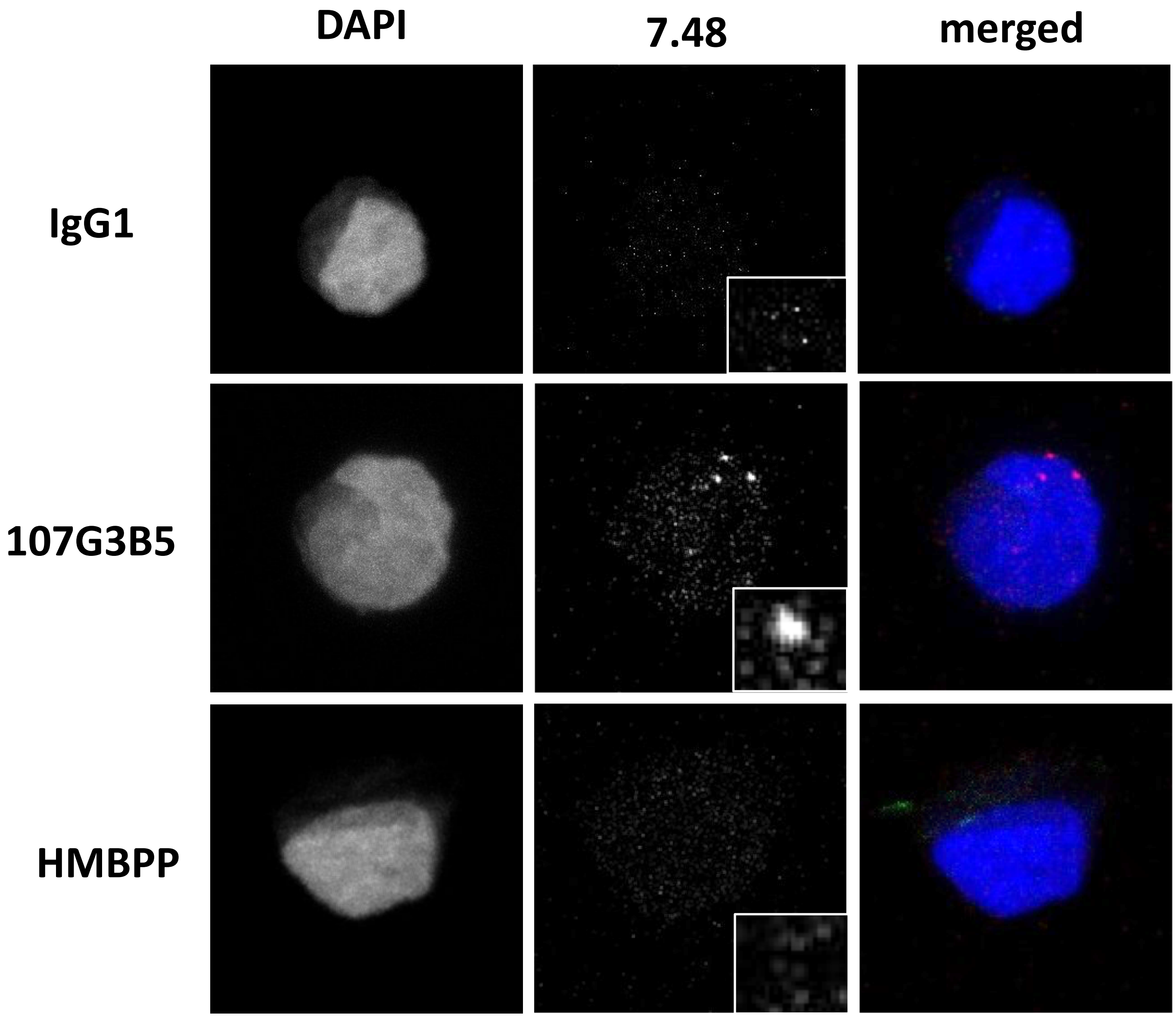


**Supplementary Figure 7. 107G3B5 induces clustering of BTN2A1.** Confocal microscopy of BTN2A on SUPT1 cell line with ou without previous saturation with an isotype control, 107G3B5 antibody or HMBPP. Evaluation BTN2A expression using aBTN2A 7.48 clone. In merged images, DAPI is represented in blue and BTN2A in red.

| | All patients<br>N = 28 (100) |
|---|---|
| Age, years | |
| Median [range] | 48 [23-80] |
| >60 | 8 (28.6) |
| Gender | |
| Male | 14 (50) |
| Female | 14 (50) |
| WBC, x10$^9$/L | |
| Median [range] | 76.1 [0.1-347] |
| High WBC count | 19 (67.8) |
| FAB subtype | |
| B | 19 (67.9) |
| B Ph- | 10 (52.6) |
| B Ph+ | 9 (47.4) |
| T | 9 (32.1) |
| PB blasts, % | |
| Mean (SD) | 84.3 (9.6) |
| Induction | 28 (100) |
| CR after induction | 100 (100) |
| Allo-SCT | 14 (50) |

**Supplementary Table 1. Baseline patient and disease characteristics.** Allo-SCT: allogenic hematopoietic stem cell transplantation in 1st complete remission; CR : complete remission; High WBC count : ≥ 30 *109/L for B lineage and ≥ 100*109 for T lineage;
Ph-: Philadelphia chromosome-negative ALL; Ph+: Philadelphia chromosome-positive ALL; PB : peripheral blood; WBC: white blood cells.
